# Awake and hungry: artificial light at night disrupts behaviour and reproductive ecology in a wild migratory bird

**DOI:** 10.1101/2025.06.10.658843

**Authors:** Juliette Champenois, Sayuri Diaz-Palma, Irene Di Lecce, Mariusz Cichoń, Lars Gustafsson, Joanna Sudyka

**Affiliations:** Institute of Environmental Sciences, Faculty of Biology, Jagiellonian University, Poland; Doctoral School of Exact and Natural Sciences, Jagiellonian University, Kraków, Poland; Institute of Evolutionary Biology, Biological and Chemical Research Centre, Faculty of Biology, University of Warsaw, Poland; Department of Animal Ecology/Ecology and Genetics, Evolutionary Biology Centre, Uppsala University, Sweden

**Keywords:** ALAN, light pollution, urbanization, migratory birds, begging behaviour, parental care, activity onset, activity offset, circadian rhythms

## Abstract

Artificial light at night (ALAN), a growing consequence of global urbanisation, disrupts natural light cues that regulate biological rhythms across taxa. However, the behavioural pathways linking ALAN exposure to broader ecological impacts remain poorly understood. Here, we experimentally introduced ALAN into nestboxes of a long-distance migratory bird, the collared flycatcher (*Ficedula albicollis*), breeding in Gotland, Sweden, to assess ALAN effects on development and reproductive ecology. Nestlings were exposed to ALAN from two days post-hatching until fledging, and we video-recorded parental and nestling activity over 24 hours on day 8 post-hatching. From a high-resolution behavioural dataset (32,100 nestling and 3,709 parental events), we found that ALAN-exposed nestlings begged more frequently and for longer at night compared to dark controls, revealing disrupted nocturnal behaviour. These effects cascaded to parental care: both females and males in ALAN nests extended their daily activity, initiating feeding earlier and terminating later, but traded off this extension by reducing their hourly feeding counts compared to parents in dark conditions. Consequently, nestlings under ALAN fledged at an older age and had a lower fledging rate after day eight post-hatching, despite no difference in total fledging numbers compared to controls. Our findings provide the first comprehensive experimental evidence that ALAN alters the behaviour of both parents alongside their offspring and reduces reproductive success in a long-distance migrant, a species group that increasingly encounters light pollution during nocturnal migration. ALAN, though directly affecting mainly nestlings and females in nestboxes, triggered socially-mediated responses that altered the circadian behaviour of entire bird families. This study underscores the need to consider behavioural disruption as a critical mechanism underlying the ecological impacts of ALAN in natural populations.

Summary figure: Cascading effects of artificial light at night, an aspect of human-induced light pollution, on the behaviour and reproductive ecology of collared flycatchers

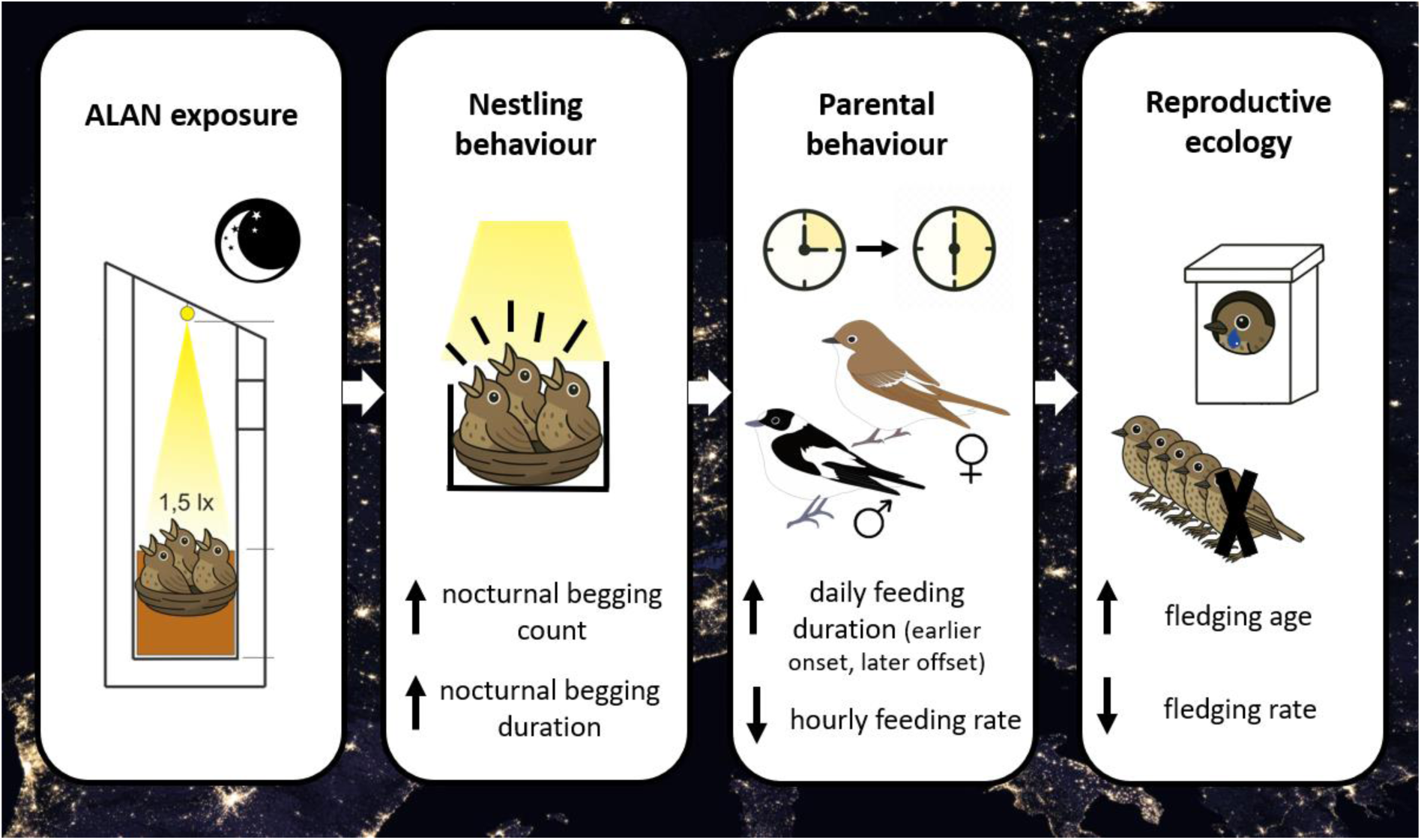

## Introduction

The timing of key life events, such as development, reproduction and migration, is critical for Darwinian fitness (Helm et al., 2017) and relies on aligning biological rhythms with consistent environmental cues, like the daily light-dark cycle. Across the animal kingdom, light universally entrains the circadian clock. In vertebrates, this biological system involves the retinae, pineal gland, and hypothalamus (Buhr & Takahashi, 2013), and acts by synchronising melatonin (sleep-regulating hormone) production to modulate physiology and behaviour (Grunst & Grunst, 2023; Helm et al., 2024). Given the critical role of light in regulating biological rhythms, anthropogenic light pollution, particularly artificial light at night (ALAN), has attracted considerable attention in behavioural research across taxa (Bedrosian et al., 2011; Owens & Lewis, 2018; Pulgar et al., 2019; Touzot et al., 2020). However, most evidence comes from controlled or captive settings, and causal links in ecologically relevant, wild environments remain poorly understood. There is therefore a pressing need for experimental ALAN manipulations in wild populations to uncover the underlying behavioural mechanisms and assess impacts on life-history traits and fitness (Sanders et al., 2021; Thoré et al., 2024).

Birds are particularly well-suited for such studies on the effects of ALAN due to their widespread distribution across diverse habitats, many of which are affected by ALAN, posing a pervasive and ecologically relevant stressor. As diurnal animals, birds rely heavily on natural light cues to rhythm daily activities, including foraging, mating, and nesting (Dawson et al., 2001). The avian circadian clock is susceptible to light, particularly via the melatonin pathway (Cassone, 2014; Gwinner & Brandstatter, 2001; Wang et al., 2012). The impact of ALAN on avian behaviour has been widely documented, with particular attention to circannual cycles linked to migration. Many diurnal migratory birds travel nocturnally (Gwinner, 1996; Rattenborg et al., 2004) and are thus exposed to prolonged artificial light, which can act as an ecological trap by disrupting navigation and concentrating birds in unsuitable habitats (Burt et al., 2023; Horton et al., 2023; Van Doren et al., 2017). ALAN also alters avian phenology by advancing laying dates (De Jong et al., 2015; Kempenaers et al., 2010) and increasing extra-pair mating (Kempenaers et al., 2010), both with potential consequences for mating dynamics. Beyond migratory and reproductive shifts, ALAN disrupts sleep patterns (Aulsebrook et al., 2018; Raap et al., 2015; Ren et al., 2022), which was linked to impaired cognitive performance (Aulsebrook et al., 2021). In addition, it is well documented that ALAN induced earlier activity onset (Beaugeard et al., 2024; Da Silva et al., 2016; Spoelstra et al., 2018), advanced vocalisations (Kempenaers et al., 2010; Silva et al., 2014), and extended evening foraging (Russ et al., 2015) in birds. During the breeding period, parents extended their daily feeding activity under ALAN (Stracey et al., 2014; Wang et al., 2021). However, the evidence on ALAN effects on provisioning rates is limited only to females, showing that the rates can be either reduced (Injaian et al., 2021) or increased (Titulaer et al., 2012).

While behavioural shifts under ALAN are well described, impacts on reproductive success remain less clear. No consistent effects were found on clutch size, hatching, or fledging success (Beaugeard et al., 2024; De Jong et al., 2015), though some species-specific effects emerged. For instance, barn swallows (*Hirundo rustica*) fledged more offspring from first broods under ALAN (Wang et al., 2021), and nestling mass in great tits (*Parus major*) varied with proximity to light (De Jong et al., 2015). In great tits, nestlings exposed to ALAN begged more at night (Raap, Pinxten, et al., 2016), thereby increasing their net energy use, resulting in impaired body mass gain (Raap, Casasole, et al., 2016). However, most studies focus on resident species, overlooking migratory birds, which are under severe time pressure to optimize the timing of migrations and coordinate them with the reproductive season in a way to maximize their fitness (Bradshaw & Holzapfel, 2007; Dawson et al., 2001; Gwinner, 1996). Addressing this gap is urgent to fully understand how ALAN reshapes behaviour, development, and survival in wild populations.

To address these challenges, we experimentally introduced ALAN into nestboxes during the breeding season of a free-ranging migratory songbird, the collared flycatcher (*Ficedula albicollis*). Using high-resolution video recordings, we were able to quantify the behavioural response to ALAN of both offspring and parents. The conspicuous sexual dimorphism in flycatchers allowed us to investigate sex-specific sensitivity to ALAN in parental behaviour (Fresneau et al., 2024; Lucass et al., 2016; Titulaer et al., 2012). To understand how ALAN alters parent-offspring behavioural dynamics, we quantified: (i) nestling begging behaviour under direct influence of ALAN, (ii) the relationship between nestling begging and parental feeding behaviours, (iii) sex-specific parental activity, including onset, offset and feeding rates and (iv) how feeding effort modulated ALAN’s impact on reproductive ecology, specifically nestling development, body condition, fledging success, and nesting period duration (age at fledging). We expected that ALAN would increase begging at night and reduce it during the day compared to controls, in terms of the number of begging events and average begging duration (Raap, Pinxten, et al., 2016). Since nestling begging behaviour and feeding by parents are positively correlated (Fresneau & Müller, 2019), we predicted that during the day, the reduced begging in ALAN-exposed nestlings would lead to diminished feeding from their parents. We hypothesized that males and females would alter their feeding behaviour under ALAN, without predicting the sex-specific direction of change [evidence limited to females so far (Injaian et al., 2021; Titulaer et al., 2012)]. However, we expected both parents to extend their daily activity periods relative to controls (Stracey et al., 2014; Wang et al., 2021). Finally, we hypothesized that these behavioural responses to ALAN would collectively reduce fitness proxies in exposed nestlings compared to controls. Specifically, we expected ALAN to impair nestling growth (measured as wing length and body mass gain), lower body condition at day 12 after hatching, increase the fledging age (a trait negatively associated with survival prospects; (Martin, 1995; Martin et al., 2018), and reduce overall and late stage (from day 8 post-hatching when the recordings took place) fledging success.

## Materials and methods

### Study species and area

The study was conducted across three breeding seasons (2022-2024) in a nestbox-breeding population of collared flycatchers (*Ficedula albicollis*) established 40 years ago on Gotland, Sweden (57°4’36’’N 18°19’27’’E, Fig. 1b). These small (average body mass 14 g), hole-nesting passerine birds reproduce in deciduous and mixed forests of Eastern, Central and South Europe and South-western Asia. Males arrive at the breeding site in May, around one week before females and all birds migrate in mid to late August to Southern Africa for their wintering grounds (Part & Gustafsson, 1989). Females lay one clutch of eggs per season, with a median size of six. Incubation in the Gotland population lasts approximately 12 days, both parents feed nestlings, and nestlings fledge at the age of ca. 16 days. Our study took place in eleven spatially distinct sites (Fig. 1b), with varying nestbox density per ha. The median nestbox density across the core areas of six sites was 6.4/ha (range: 4.5-8.7). The wooden nestboxes (Fig. 1a) are located ca. 25-50 meters apart, and the entrance hole is located on average at 120 cm (SD ± 14 cm) above the ground. The vegetation varies across the sites, but is mainly dominated by oaks (*Quercus robur*), ash (*Fraxinus excelsior*) and poplar (*Populus spp*) with an understory of hazel (*Corylus avellana*). Some sites are located within a coniferous forest dominated by pine (*Pinus sylvestris L.*), with birch (*Betula spp.*), or a forest-meadow with more sparse trees, formerly used as pastures. Gotland’s climate is semi-continental with moderate rainfall during spring and summer (Ebert et al., 2016). We used temperature data loggers (Maxim Integrated DS1921G; range: −40 °C to +85 °C; precision: ± 1 °C; resolution: 0.5 °C), placed at ~120 cm above the ground in a protective net shading from rain and direct sunlight at one of our study sites to measure average ambient temperatures. These were 13°C in May and 15°C in June (2022 - 2023), ranging from a minimum of 3°C to a maximum of 27°C. Due to the island’s northern location, nighttime is particularly short. In May and June, the sun rose on average at 4:24 and 03:48 respectively, and set at 21:01 and 21:43 respectively.

**Figure 1 –.**
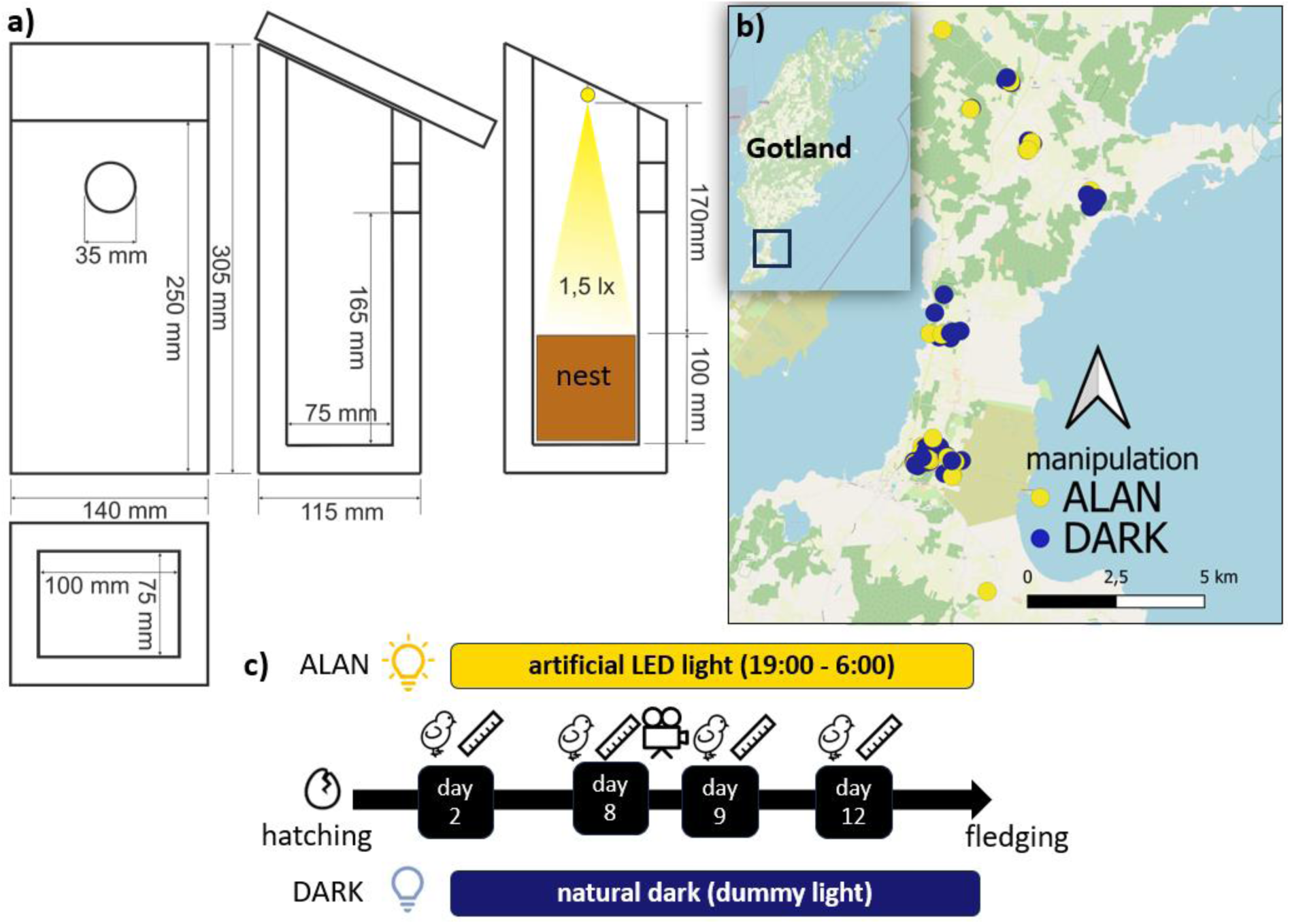
a) Nestbox dimensions with the LED light position, b) location of the study sites with ALAN and DARK nestboxes marked with dots, and c) experimental timeline, where a symbolic ruler denotes biometric measurements of nestlings, and a camera denotes the time of recordings.

### Fieldwork, light sets, cameras and video analysis

A detailed description of all procedures can be found in the Supplementary Material. In short, nestboxes were inspected every four days from the end of April to record all breeding attempts. After identifying an active flycatcher nest, we started individual nest monitoring. We paired two nests with matching hatching date (HD ± 1 day) and number of hatchlings (brood size ± 1), thereby creating experimental pairs (a factor in statistical analyses), to control for environmental variables (especially day duration) and within-nest competition, and randomly assigned them to either the experimental (ALAN) or control (DARK) group (Fig. 1c). In the ALAN group, two days after hatching, we installed a white LED light diode producing at the average level of nest cup 1.5 lx (Fig. 1a), which is a realistic magnitude of light pollution in anthropogenically altered habitats (Sanders et al., 2021). While direct exposure to light pollution in nest cavities is limited, ALAN can enter through entrance holes or diffuse via any breaks in cavity walls. Our in-box illumination scheme simulates this exposure, reflecting real-world, ecologically relevant scenarios in urban and peri-urban environments. Moreover, by focusing on direct behavioural and physiological responses, we can better understand how these migratory birds cope with or adapt to ALAN exposure both within the nest cavity and outside. The light diode was controlled by Arduino and programmed to switch on at 19h00 and off at 06h00 (Fig. S1). In the DARK group, a dummy light was placed instead (inactive diode). To record the behaviour inside the nest, small spy cameras were magnetically fixed next to the diode on HD + 8 for ca. 24 hours. HD + 8 was selected because around this time the energetic cost of rearing passerine nestlings (Lemon, 1993; Sudyka et al., 2016) and parental feeding rates (ten Cate, 1982) are the greatest. We used eight KA9 cameras by kamREC in 2022 and eight A10 cameras by NOBITECH SC in 2023 (Fig. S2). We also collected biometric measurements such as body mass, tarsus and wing length (Fig. 1c, Table S1), uniquely ringed and blood sampled each nestling via brachial vein puncture for molecular sexing (see Supplementary Material). After confirming fledging (checks started on HD + 15), the light sets were removed and dead nestlings (if any) were identified by their ring number to determine fledging success. A total of 45 nests were recorded and analysed in 2022 and 2023. 21 nests in the 2024 season were added, totalling 66 for the analyses of first and last parental visit (Table S2).

We established an ethogram (Table S3) by carefully screening (blindly regarding the treatment) one hour of night (between 01:00 and 02:00) and one hour of day (between 09:00 and 10:00) activity per nest. While we recorded ca. 24h of activity per nest, we decided to reduce the detailed analysis to just two hours because of the number and duration of the recordings, and because one hour represents overall activity within the nest (Supplementary Material). We obtained behavioural data from the videos using BORIS (Behavioral Observation Research Interactive Software), a cross-platform tool for coding and analysing behavioural data from videos (Friard & Gamba, 2016). Each behaviour type (Table S3), either a point event (single occurrence) or a state event (continuous occurrence of a certain duration), was attributed to one subject: the female, the male or the nestlings. Because nestlings were not visibly marked, individual behaviours could not be recorded, so a behaviour was noted whenever exhibited by at least one nestling. Nestling behaviours encompassed begging, moving, scratching, and wiggling. Adult behaviours encompassed entering the nestbox, feeding, cleaning, moving, scratching, sitting, wiggling, sleeping and allofeeding. Across all nests, we recorded 32,100 behaviours in nestlings (17,457 at night and 14,643 during the day) and 3,709 in parents (401 at night and 3,308 during the day). Subsequently, we focused on parameters related to begging and feeding behaviours, because they are key indicators of parent-offspring interactions and energy allocation strategies. Relating to nestling behaviour, for each nest we quantified: total begging count (number of begging events summarised for night and day), begging during day (hourly number of begging events during the day), begging at night (hourly number of begging events during the night), and begging duration (average duration of a begging event in seconds). Relating to parental behaviour, for each nest we quantified: feeding count (total hourly number of feeding events), male feeding count (hourly number of feeding events by male), female feeding count (hourly number of feeding events by female), parent visit duration (average duration of parental visit during the day in seconds), and entering nest count (hourly number of parental visit events during the day, including feeding events). In one ALAN nest, we did not have the daily hour recording and in two nests (one ALAN and one DARK), we did not have the nightly hour recording due to mistakes in the motion detection setting. We observed strong correlations among the behavioural parameters (Fig. S5). Parental daily activity offset was assessed from video recordings starting on the evening of camera installation (HD + 8). Last visits were defined as the parents’ final departure before evening inactivity, analysed separately by sex. In seven cases where females stayed overnight, the time before settling for the night was used. First visits were recorded the next morning (HD + 9) as the parents’ initial appearance, or, for those same females, their reappearance after first leaving. The times of the last and first visits were converted to minutes after sunset and after sunrise respectively.

### Statistical analysis

We tested the effects of ALAN on nestling and parental behaviour, and reproductive outcomes using a series of generalized linear mixed models (GLMMs) and linear mixed models (LMMs) in R version 4.2.2 (R Core Team, 2022), with AIC-based model selection via the *dredge*() function in MuMIn (Bartoń, 2022). Random effects structures included study site, experimental pair, and nest ID (nested in pair), unless noted otherwise. The *set of fixed effects* in all models included manipulation (ALAN vs DARK), year of study, brood size at day 8, day 8 date (as days from April 1^st^, timing in the breeding season), average nestling body mass at day 8, and proportion of male offspring [sex-specific nutritional requirements of nestlings can influence parental provisioning and fitness outcomes (Sheldon et al., 1998)]. All models retained a focal interaction relevant to each hypothesis, and we used the *fixed* condition in *dredge()* to ensure it was always retained as a variable of interest.

#### Model 1: Circadian begging patterns

We tested hourly begging counts (GLMM, negative binomial, with the package glmmTMB (Brooks et al., 2017) to account for zero-inflation, model 1a) and begging duration (LMM, response natural log-transformed, model 1b); the models additionally included time (day or night) and the focal interaction of manipulation × time as fixed effects.

#### Model 2: Parent-offspring communication

To test whether the begging duration of the nestlings was stimulated by the average time parents spent in the nest, we modelled natural log-transformed begging duration per hour during daytime (LMM) with parent visit duration in addition to the fixed effects, and included the focal interaction manipulation × parent visit duration.

#### Models 3-4: Parental activity timing and feeding patterns

We assessed activity offset (last visit, model 3a) and onset (first visit, model 3b) in two separate LMMs. Fixed effects additionally included parental sex, and the focal interaction was manipulation × sex. Since male and female feeding counts were not correlated (Fig. S5), we fitted separate negative binomial glmmTMB models for female (model 4a) and male feeding counts (model 4b) during the day without the random effect nest ID. An additional fixed effect was the nestling begging counts during the day, and the focal interaction manipulation × begging counts.

#### Models 5–10: Development and fitness proxies

In addition to *the set of fixed effects* described above, total feeding counts (male + female), and the focal interaction manipulation × total feeding count were also included. These models did not include nest ID in the random effect structure. Model 5) on wing growth (day 8 to 9, LMM) additionally included wing length at day 8, instead of body mass day 8, and as random effects the exact time between measurements in hours (as these ranged from 24 to 30 hours), and observer ID (the person who measured the structural size each day, due to inter-observer variation inherent to this type of measurement). Model 6) on body mass gain (day 8 to 9, LMM) included the exact time between measurements in hours. Model 7) on nestling body condition [measured as Scale Mass Index (SMI) on day 12, (Peig & Green, 2009); LMM] additionally included ectoparasite presence at day 12 (which could negatively influence body condition and fledging), and the random effect observer ID. Model 8) on fledgling number, a universal measure of avian reproductive success (LMM), included ectoparasite presence and SMI. Model 9) on fledging ratio, as a number of fledged offspring over a number of offspring in the nest at day 8, when the recording took place (GLMM, binomial), was weighted by brood size at day 8. Model 10) on age at fledging, an indicator of parental investment [more extended nesting periods may enhance survival through increased parental care, see Bowers et al., (2013); Martin et al., (2018); LMM] included ectoparasite presence, SMI and wing length at day 12 (more developed wing would possibly result in faster fledging). In models 7), 8) and 10, we focused on day 12 biometrics to better represent the complete brood trajectory (from hatching till closest to fledging) and to align with SMI thus we included hatchling number instead of brood size d8 and day 12 date (instead of day 8 date) and did not include average nestling body mass at day 8.

To identify the most parsimonious model in each case, we used AICc-based model averaging (MuMIn::dredge). Predictors from models with ΔAICc < 2 (Burnham & Anderson, 2002) were retained and refitted in a single linear mixed-effects model to obtain parameter estimates, random effect variance, VIF scores, and R^2^ (Bartoń, 2022; Nakagawa & Schielzeth, 2013). Due to singular fit issues during model selection, random effects with (near-)zero variance were removed prior to *dredge()*. VIF values for final models were <3 (except for terms in interactions). Model fit was assessed using KS tests, dispersion checks, and DHARMa residual diagnostics (Hartig, 2024). Outliers were tested by rerunning models on adjusted datasets; as results were qualitatively unchanged, all data points were retained. Pairwise group differences were tested using the package emmeans (Lenth, 2017). To compare the full set of circadian nestling and parental behaviours (ALAN vs DARK, separately for day and night, Figs S3-4), and the counts of nestling behaviours outside of begging (ALAN vs DARK, Fig. S3), we used *Χ^2^* tests.

## Results

### Nestling begging behaviour

ALAN significantly disrupted nestling circadian begging activity, with a strong interaction between treatment and time of day for begging occurrence (z = 8.19, p < 0.0001; Table 1a, S4a; Fig. 2a) and duration (t = 3.03, p = 0.002; Table 1b, S4b; Fig. 2b). Begging occurrence (−0.23 ± 0.10; z = 2.39, p = 0.017), but not duration (−0.01 ± 0.04; t = −0.30, p =0.765), was markedly reduced under exposure to ALAN. Unlike control nestlings, which showed near-complete cessation of begging at night (Fig. 2a), ALAN-exposed nestlings continued to beg after dark at elevated levels and they did so for longer periods of time (Fig. 2b). Broods with on average heavier nestlings begged less often (−0.14 ± 0.04; z = −3.30, p = 0.0009).

**Figure 2 -.**
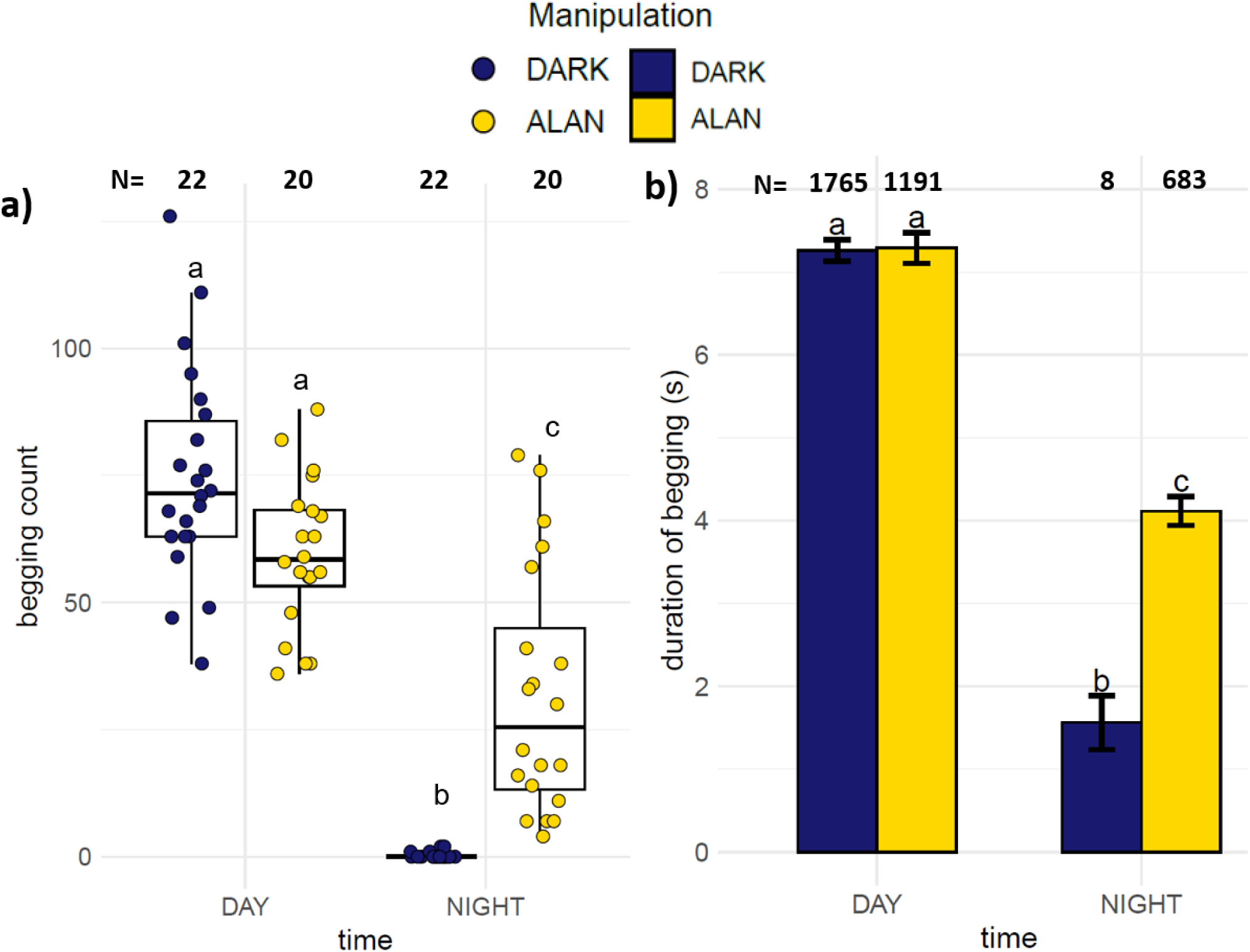
a) Begging occurrence (count per hour per nest) of nestlings under DARK (dark blue) and ALAN (yellow) for day and nighttime. The boxes represent the median with IQR and the whiskers 1.5 IQR plotted on raw values. b) Duration of begging recorded in nestlings under DARK (dark blue) and ALAN (yellow) manipulation during the day (white background), and at night (grey background). Mean and standard errors are shown. Letters above indicate significantly different groups according to post hoc tests.

**Table 1 -.** Models examining variation in circadian hourly a) begging count and b) begging duration. (natural log-transformed for normality of the model residual distribution) as the dependent variables in nestling collared flycatchers. Predictor variables were selected based on model averaging across all models with ΔAICc <2, where Manipulation:Time was always retained as a *fixed* factor of interest. Significant differences (P < 0.05) are indicated in bold. Marginal (R^2^m) and conditional (R^2^c) R-squared are shown.

|  | Estimate | SE | z | p |
| --- | --- | --- | --- | --- |
| (Intercept) | 6.92 | 1.57 | 4.40 | **<0.0001** |
| Manipulation (ALAN) | -0.23 | 0.10 | -2.39 | **0.017** |
| Time (Night) | -4.64 | 0.46 | -10.06 | **<0.0001** |
| BroodSizeD8 | 0.03 | 0.04 | 0.86 | 0.390 |
| Day8Date | -0.01 | 0.02 | -0.76 | 0.450 |
| ProportionMaleOffspring | -0.16 | 0.21 | -0.76 | 0.444 |
| AverageNestlingMassD8 | -0.14 | 0.04 | -3.30 | **0.0009** |
| Manipulation (ALAN):Time (Night) | 3.92 | 0.48 | 8.19 | **<0.0001** |
| *Random effects* | *Variance* |  |  |  |
| *Nest id:Pair* | 0 (removed) |  |  |  |
| *Pair* | 0 (removed) |  |  |  |
| *Site* | 0 (removed) |  |  |  |
| $R^2_m$ / $R^2_c$ | 0.958 / 0.958 | | | |

**b. Model 1b: circadian begging duration, LMM gaussian** N = 3484 observations in 43 nests
|  | Estimate | SE | t | p |
| --- | --- | --- | --- | --- |
| (Intercept) | 1.41 | 0.11 | 13.23 | **<0.0001** |
| Manipulation (ALAN) | -0.01 | 0.04 | -0.30 | 0.765 |
| Time (Night) | -1.41 | 0.28 | -4.99 | **<0.0001** |
| BroodSizeD8 | 0.05 | 0.02 | 2.74 | 0.009 |
| Manipulation (ALAN):Time (Night) | 0.87 | 0.29 | 3.03 | **0.002** |
| *Random effects* | *Variance* |  |  |  |
| *Nest id:Pair* | 0.007 |  |  |  |
| *Pair* | 0.003 |  |  |  |
| *Site* | 0 (removed) |  |  |  |
| $R^2_m$ / $R^2_c$ | 0.078 / 0.091 | | | |

When examining daytime begging duration, there was a statistical tendency for longer parental visits to be associated with longer begging bouts (0.007 ± 0.004; t = 1.84, p = 0.074, Table S5). However, ALAN exposure alone had no significant effect (−0.02 ± 0.05; t = −0.29, p = 0.771) and did not interact with visit duration (0.011 ± 0.007, t = 1.34, p = 0.188), suggesting that ALAN did not alter the relationship between parental presence and nestling begging behaviour during the day. The most substantial positive contributor to begging duration was brood size (0.09 ± 0.02, t = 3.90, p = 0.0004).

### Parental behaviour: offset and onset of daily activity and feeding behaviour

Exposure to ALAN significantly extended parental activity periods. Parents exposed to ALAN delayed their last visit by 23 ± 4.5 minutes (t = 5.09, p < 0.0001, Table 2a), and advanced their first visit the following morning by 17 ± 3.7 minutes before sunrise (t = −4.72, p < 0.0001, Table 2b) compared to controls, resulting in an overall increase of approximately 40 minutes in daily activity duration (Fig. 3). Males ceased their daily activity earlier than females, with last visits occurring 9 ± 3.6 minutes earlier on average (t = −2.33, p = 0.023, Table 2a; Fig. S6a). Additionally, parental activity timing was influenced by both phenology and brood traits: later hatching dates were associated with slightly earlier last visits (−3 ± 1.0 minutes, p = 0.018, Table 2a; Fig. S6c), and higher average nestling body mass on day 8 predicted later final visits (4 ± 1.9, p = 0.025, Table 2a; Fig. S6b), suggesting more intense parental care in higher-quality broods. Nests with more male nestlings were visited earlier in the morning (−17 ± 8.6 minutes, p = 0.046, Table 2b; Fig. S6d). However, the interaction between treatment and sex was not significant for either last visit (−1.24 ± 7.41, t = −0.17, p = 0.867) or first visit (2.13 ± 3.61, t = 0.60, p = 0.554), indicating that the effects of ALAN on activity timing did not differ between males and females.

**Figure 3 -.**
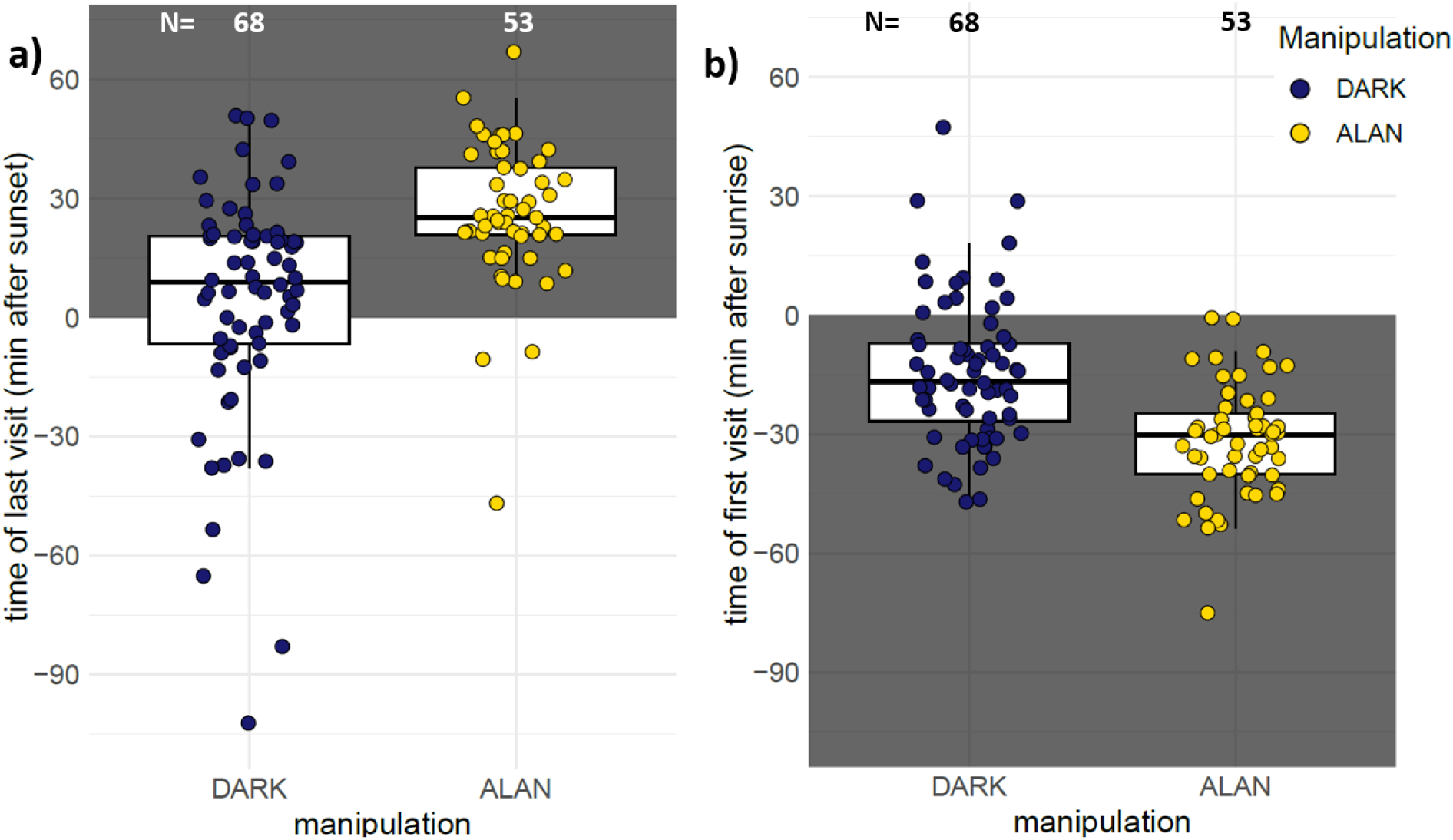
Boxplot of the timing of a) last parental visit on the 8^th^ day after hatching (left panel) and b) first visit on the 9^th^ day after hatching, under ALAN (yellow) or DARK (dark blue) conditions. A white background indicates daytime (before sunset and after sunrise), while nighttime (after sunset and before sunrise) is indicated by a grey background. The boxes represent the median with IQR, and the whiskers 1.5 IQR. Plotted on raw values.

**Table 2 -.** Models examining variation in the offset and onset of parental activity with last visit as minutes after sunset and first visit as minutes after sunrise. Predictor variables were selected based on model averaging across all models with ΔAICc <2, where Manipulation:Sex was always retained as a fixed factor of interest. For last visit the best model is presented (since just one model’s ΔAICc <2). Significant differences (P < 0.05) are indicated in bold. Marginal (R^2^m) and conditional (R^2^c) R-squared are shown. Since the focal interaction Manipulation:Sex was non-significant in both models (−1.24 ± 7.41, t = −0.17, p = 0.867 for last visit and 2.13 ± 3.61, t = 0.60, p = 0.554 for first visit), we removed those for the correctness of main factor interpretation in the final models.

|  | Estimate | SE | t | p |
| --- | --- | --- | --- | --- |
| (Intercept) | 130.84 | 84.95 | 1.54 | 0.135 |
| Manipulation (ALAN) | 22.90 | 4.50 | 5.09 | **<0.0001** |
| ParentSex (Male) | -8.51 | 3.65 | -2.33 | **0.023** |
| Day8Date | -2.59 | 1.03 | -2.51 | **0.018** |
| BroodSizeD8 | 2.54 | 1.99 | 1.28 | 0.209 |
| ProportionMaleOffspring | -1.82 | 10.38 | -0.18 | 0.862 |
| AverageNestlingMassD8 | 4.37 | 1.89 | 2.32 | **0.025** |
| Year (2023) | 1.68 | 6.94 | 0.24 | 0.811 |
| Year (2024) | 2.20 | 8.17 | 0.27 | 0.790 |
| *Random effects* | *Variance* |  |  |  |
| *Nest id:Pair* | 84.581 |  |  |  |
| *Pair* | 24.140 |  |  |  |
| *Site* | 6.525 |  |  |  |
| **R<sup>2</sup>m / R<sup>2</sup>c** | 0.320/ 0.476 |  |  |  |

b. Model 3b: First visit, LMM gaussian N = 121 observations in 63 nests
|  | Estimate | SE | t | p |
| --- | --- | --- | --- | --- |
| (Intercept) | -72.75 | 65.00 | 1.12 | 0.268 |
| Manipulation (ALAN) | -17.37 | 3.68 | -4.72 | **<0.0001** |
| ParentSex (Male) | 0.74 | 1.78 | 0.42 | 0.678 |
| Day8Date | 1.44 | 0.79 | 1.82 | 0.074 |
| BroodSizeD8 | -1.67 | 1.56 | -1.07 | 0.289 |
| ProportionMaleOffspring | -16.85 | 8.26 | -2.04 | **0.046** |
| AverageNestlingMassD8 | -2.73 | 1.47 | -1.86 | 0.068 |
| Year (2023) | 5.63 | 5.31 | 1.06 | 0.294 |
| Year (2024) | 2.11 | 6.26 | 0.34 | 0.737 |
| *Random effects* | *Variance* |  |  |  |
| *Nest id:Pair* | 153.250 |  |  |  |
| *Pair* | 0 (removed) |  |  |  |
| *Site* | 0 (removed) |  |  |  |
| **R<sup>2</sup>m / R<sup>2</sup>c** | 0.336/ 0.749 |  |  |  |

Sex-specific models on parental feeding rates confirmed a significant negative effect of ALAN on provisioning in both females (−0.92 ± 0.39; z = −2.37, p = 0.018, Table 3, Fig 4a) and males (−0.33 ± 0.15; z = −2.28, p = 0.022, Table 3, Fig. 4b).

**Figure 4 -.**
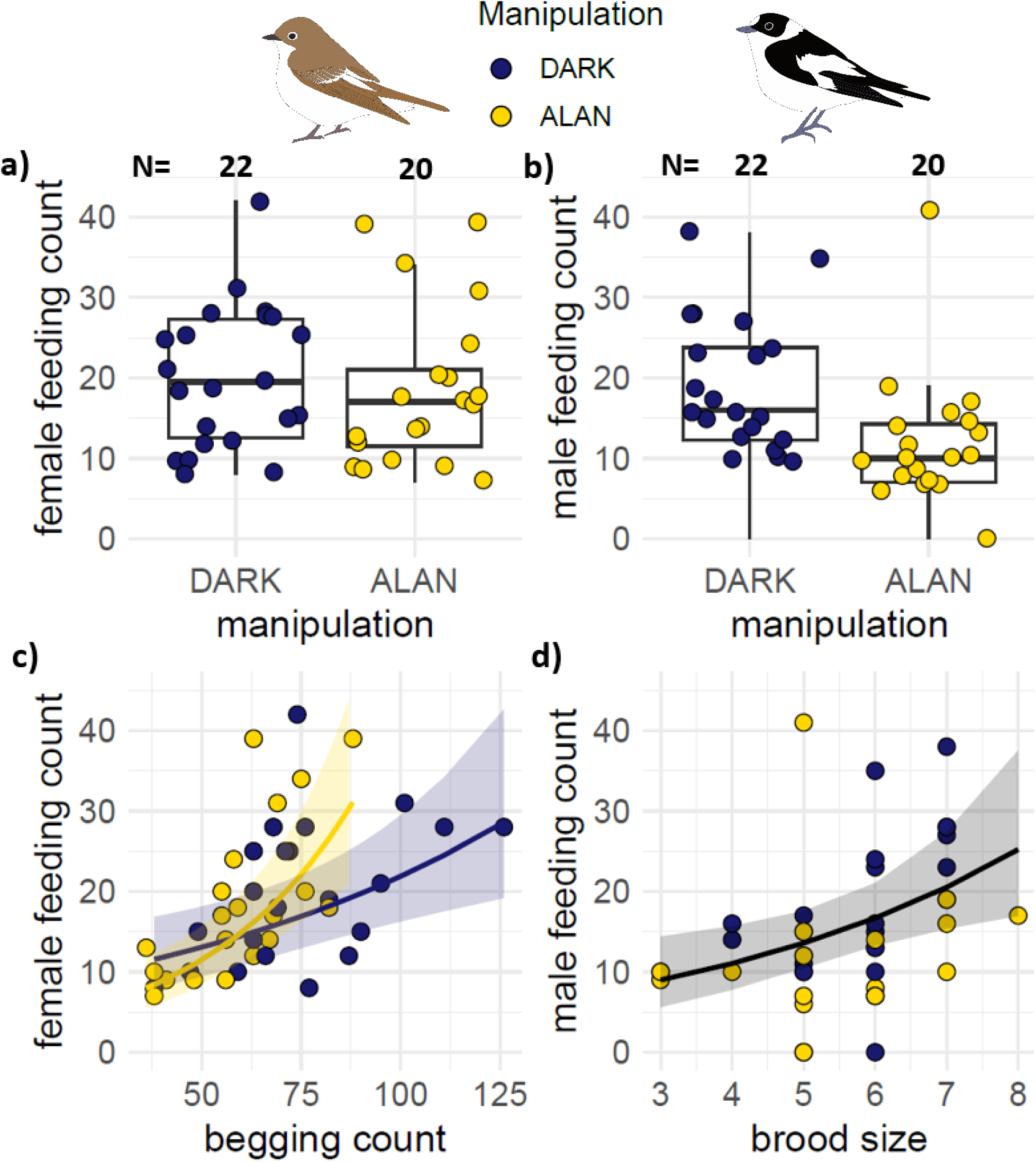
Feeding count response according to manipulation in a) females and b) males, c) females according to nestling begging count, and d) males according to brood size under ALAN (yellow) and DARK (dark blue). a), b) The boxes represent the median with IQR, and the whiskers 1.5 IQR; c), d) the lines represent model-predicted feeding counts based on negative binomial GLMM, averaging over other variables. The shaded area indicates the 95% CI of the predictions.

**Table 3 -.**
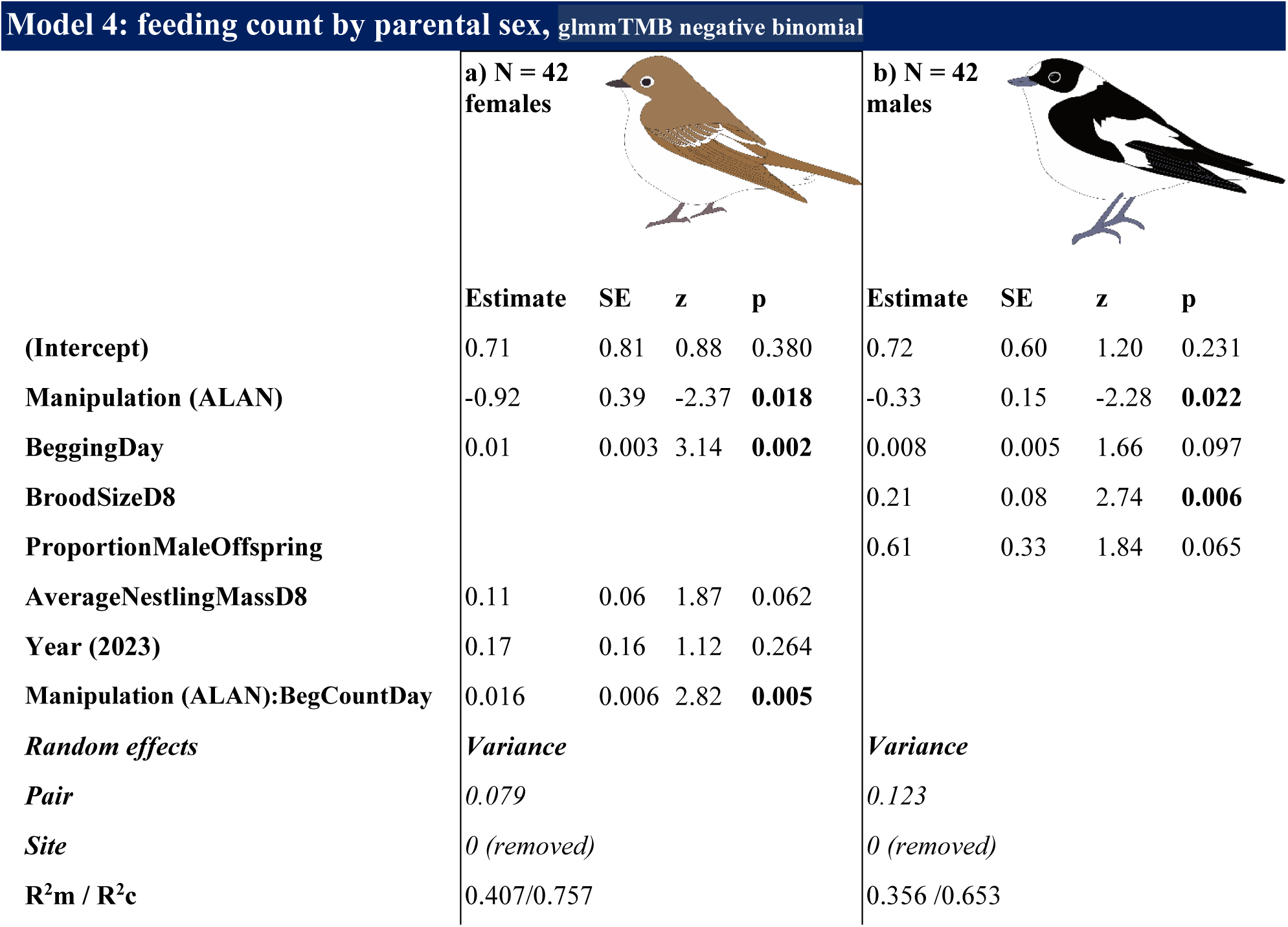
Models examining variation in parental feeding count in response to nestling begging count during the day. Predictor variables were selected based on model averaging across all models with ΔAICc <2, where Manipulation:BeggingDay was always retained as a *fixed* factor of interest. Significant differences (P < 0.05) are indicated in bold. Marginal (R^2^m) and conditional (R^2^c) R-squared are shown. Since the focal interaction Manipulation:BeggingDay in the Male Feeding Count model was non-significant (−0.008 ± 0.009, z = −0.85, p = 0.394) we removed it for the correctness of the main factor interpretation in the model.

**Model 4: feeding count by parental sex, glmmTMB negative binomial**
|  | a) N = 42 females |  |  |  | b) N = 42 males |  |  |  |
| --- | --- | --- | --- | --- | --- | --- | --- | --- |
|  | Estimate | SE | z | p | Estimate | SE | z | p |
| (Intercept) | 0.71 | 0.81 | 0.88 | 0.380 | 0.72 | 0.60 | 1.20 | 0.231 |
| Manipulation (ALAN) | -0.92 | 0.39 | -2.37 | **0.018** | -0.33 | 0.15 | -2.28 | **0.022** |
| BeggingDay | 0.01 | 0.003 | 3.14 | **0.002** | 0.008 | 0.005 | 1.66 | 0.097 |
| BroodSizeD8 |  |  |  |  | 0.21 | 0.08 | 2.74 | **0.006** |
| ProportionMaleOffspring |  |  |  |  | 0.61 | 0.33 | 1.84 | 0.065 |
| AverageNestlingMassD8 | 0.11 | 0.06 | 1.87 | 0.062 |  |  |  |  |
| Year (2023) | 0.17 | 0.16 | 1.12 | 0.264 |  |  |  |  |
| Manipulation (ALAN): BegCountDay | 0.016 | 0.006 | 2.82 | **0.005** |  |  |  |  |
| Random effects | Variance |  |  |  | Variance |  |  |  |
| Pair | 0.079 |  |  |  | 0.123 |  |  |  |
| Site | 0 (removed) |  |  |  | 0 (removed) |  |  |  |
| R<sup>2</sup><sub>m</sub> / R<sup>2</sup><sub>c</sub> | 0.407/0.757 |  |  |  | 0.356 / 0.653 |  |  |  |

Begging frequency strongly and positively influenced feeding behaviour only in females (0.01 ± 0.003; p = 0.002, Fig. 4c), and this effect was significantly amplified under ALAN (interaction: z = 2.82, p = 0.005, Table 3), indicating that light exposure reinforced female responsiveness to begging. In males, while the effect of begging frequency was only marginal (p = 0.097), brood size had a strong positive impact on feeding (0.21 ± 0.08; p = 0.006, Fig. 4d). Interestingly, there was also a trend for increased male feeding with a higher proportion of male offspring (0.61 ± 0.33; p = 0.065).

### ALAN vs reproductive ecology: nestling development, body condition, reproductive success and parental investment

Exposure to ALAN was associated with a tendency toward increased structural growth, specifically greater average wing size development (0.43 ± 0.24; t = 1.80, p = 0.094, Table S6a), although this effect was not statistically significant. In contrast, ALAN did not affect average body mass gain (0.09 ± 0.16; t = 0.54, p = 0.593, Table S6b). Feeding frequency significantly predicted wing development, with higher feeding rates linked to greater average wing gain (0.04 ± 0.02; t = 2.63, p = 0.016), but had no effect on average mass gain (p = 0.902). Brood size negatively affected wing growth, with larger broods exhibiting reduced average wing size gain (−0.89 ± 0.16; t = −5.69, p < 0.0001, Table S6a).

In the model predicting nestling body condition as day 12 SMI, ALAN exposure had no significant effect (0.01 ± 0.33, t = 0.03, p = 0.976, Table 4a), and feeding count had a significant positive effect (0.04 ± 0.02, t = 2.25, p = 0.031, Fig 5a), indicating that higher parental feeding rates were associated with improved nestling condition. The number of fledged offspring was not affected by ALAN exposure (0.12 ± 0.46, t = 0.25, p = 0.802, Table 4b), only parental feeding count had a significant positive effect (0.05 ± 0.02, t = 2.42, p = 0.021, Fig. 5b), indicating that more frequent feeding was associated with higher fledging success. The proportion of offspring fledged from day 8 was significantly reduced under ALAN exposure (−18.63 ± 5.06, z = −3.68, p = 0.0002, Table 4c). Notably, ALAN interacted significantly with parental feeding frequency (0.84 ± 0.23, z = 3.64, p = 0.0003, Fig. 5c), indicating that under artificial light, increased feeding was associated with higher fledging success. Feeding count (p = 0.023) and average nestling mass at day 8 (p = 0.0009) also positively influenced fledging proportion, but larger brood size was related to lower fledging proportion (p = 0.014, Table 4c). Nestlings under ALAN stayed in the nest longer (0.80 ± 0.33, t = 2.45, p = 0.019, Table 4d, Fig. 5d), pointing to an increased duration of parental investment. Additionally, longer wing lengths were negatively associated with fledging age (−0.17 ± 0.05, t = –3.14, p = 0.003, Fig. S7), revealing that nestlings with more developed wings left the nest sooner. The interaction between ALAN and feeding frequency was not significant in the models on body condition (0.01 ± 0.03, t = 0.49 p = 0.628), number of fledged young (−0.01 ± 0.04, t = −0.27 p = 0.791) and fledging age (−0.03 ± 0.03, t = −1.01 p = 0.321), indicating that the effect of feeding on these parameters was consistent across treatments.

**Figure 5 –.**
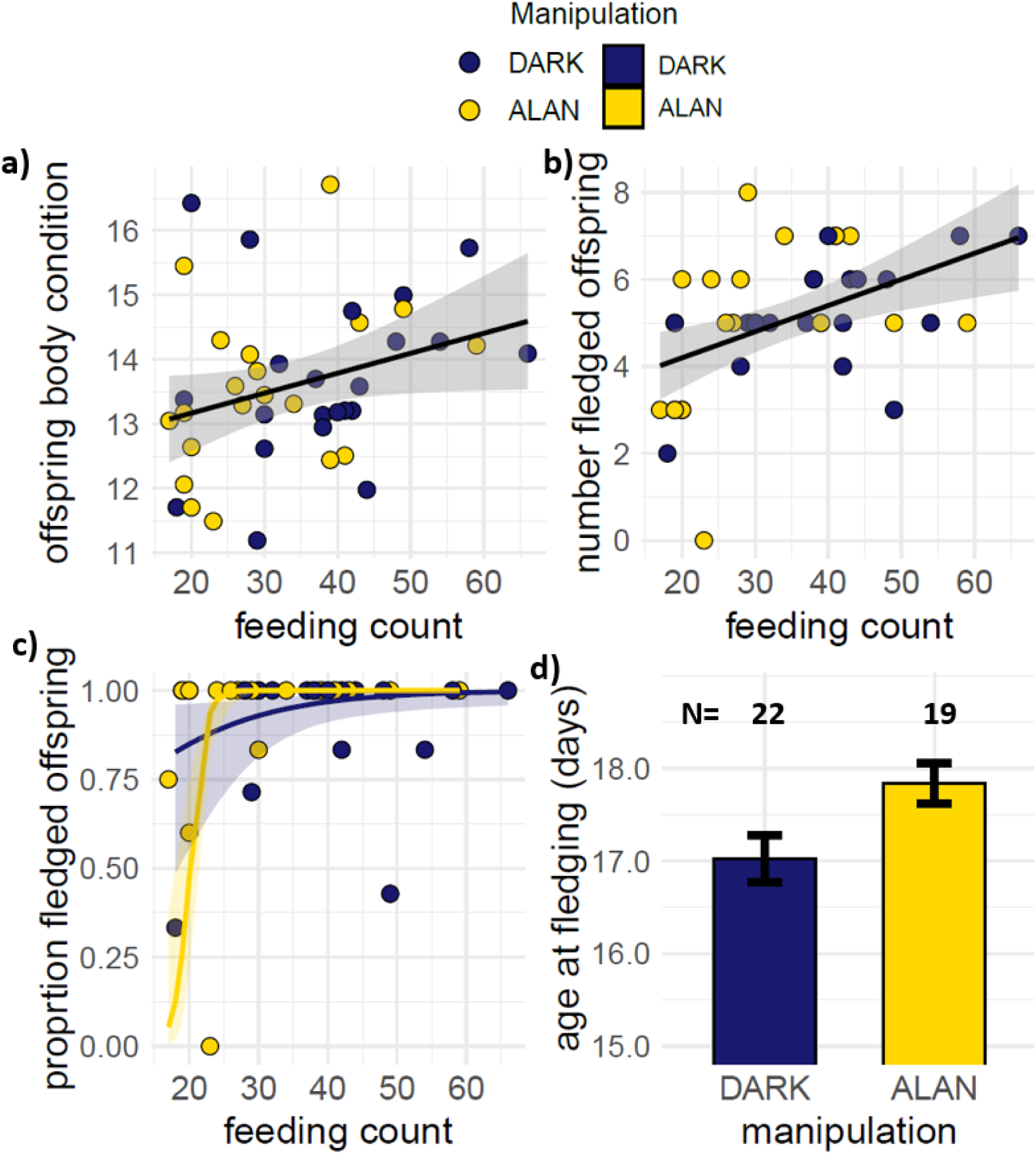
Effects of parental feeding count on a) offspring body condition as Scaled Mass Index on day 12 post-hatching, b) reproductive success as number of fledged young, c) reproductive success as proportion of offspring fledged from day 8^th^ when the feeding counts took place and d) effect of manipulation on a proxy of parental investment - age at fledging under ALAN (yellow) and DARK (dark blue) conditions. In a) and b), the lines show fitted linear relationships with shaded standard error bands, in c) the lines represent model-predicted relationships based on binomial GLMM, averaging over other variables, with 95% CI colour-shaded according to manipulation, and in d) mean and standard errors are shown.

**Table 4 -.**
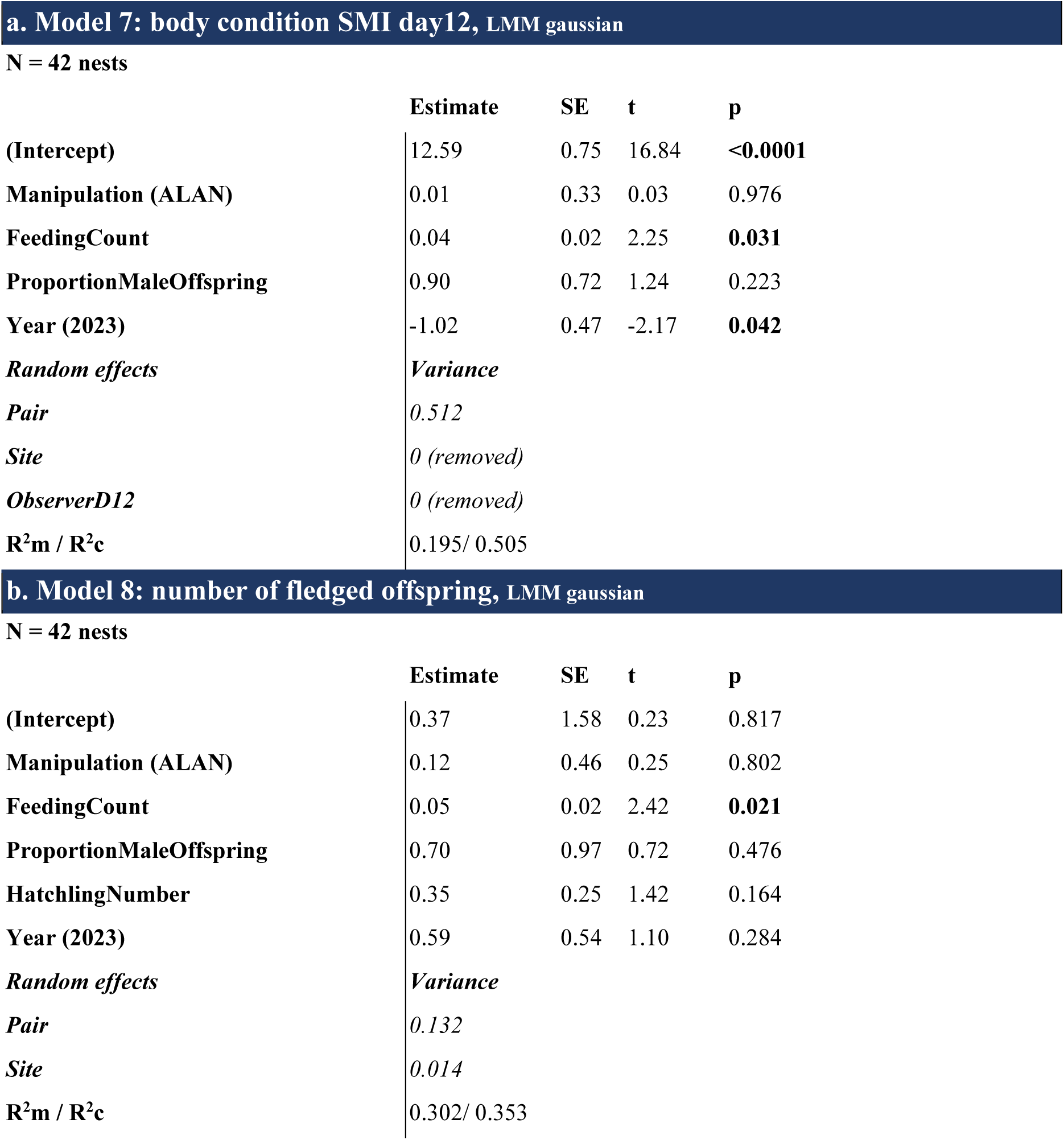
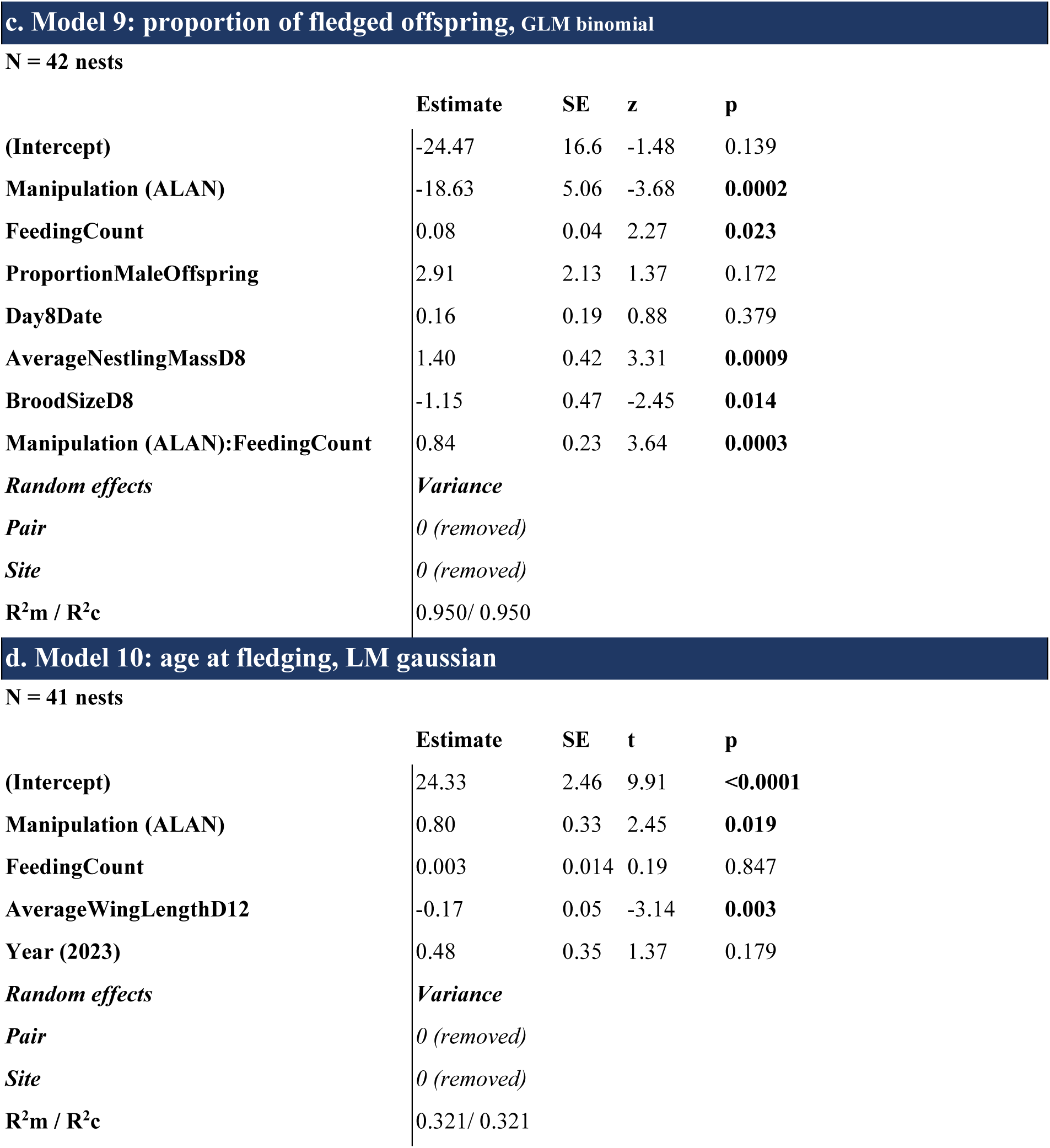
Models examining variation in a) body condition on day 12 as Scaled Mass Index, b) number of fledged offspring, c) proportion of fledged offspring from day 8 when the recording took place, d) age at fledging (days), in response to parental hourly feeding count. Predictor variables were selected based on model averaging across all models with ΔAICc <2, where Manipulation:FeedingCount was always retained as a *fixed* factor of interest. Significant differences (P < 0.05) are indicated in bold. Marginal (R^2^m) and conditional (R^2^c) R-squared are shown. Since the focal interaction Manipulation:FeedingCount in the body condition model (0.01 ± 0.03, t = 0.49 p = 0.628), number of fledged offspring model (−0.01 ± 0.04, t = −0.27 p = 0.791) and age at fledging model (−0.03 ± 0.03, t = −1.01 p = 0.321) were non-significant we removed them for the correctness of main factor interpretation in the models.

## Discussion

Our study provides the first comprehensive experimental evidence that ALAN disrupts nestling behaviour, reshapes parental care patterns, and compromises reproductive outcomes in a wild migratory bird. By leveraging continuous 24-hour behavioural monitoring of nestlings and parents, we showed that ALAN exposure leads to a fundamental breakdown in the circadian regulation of offspring begging. This nocturnal disruption triggered extended parental activity, beginning earlier and ending later, but came at the cost of reduced hourly feeding rates. Critically, these cascading behavioural changes resulted in prolonged nestling periods and reduced fledging success. Our findings reveal an underappreciated pathway through which ALAN compromises avian reproductive performance in free-living populations. Given the rapid global expansion of light pollution and the heightened sensitivity of migratory species to nocturnal environmental cues (Rebke et al., 2019), our results indicate an urgent need to integrate behavioural disruption into the ecological risk assessments of ALAN.

Circadian behaviour types differed between DARK and ALAN nests in nestlings (Fig. S3) and adults (Fig. S4). Nestlings exposed to ALAN begged more frequently and for longer durations during the night than those in the control conditions (Fig. 2, Table 1). This is consistent with previous findings in other avian species (Raap, Pinxten, et al., 2016; Wang et al., 2021). Begging behaviour is considered to convey information about the nestlings’ short- (Fresneau et al., 2018) and long-term nutritional needs (Price et al., 1996), and the rate and duration of begging increases with food deprivation (Fresneau et al., 2018; Kilner & Johnstone, 1997; Price et al., 1996). This accords with our study, where broods with heavier nestlings begged less often (Table 1a), possibly reflecting reduced need or motivation due to better nutritional status. Despite the absence of parental feeding at night, nestlings exposed to ALAN begged more, possibly due to the circadian clock mismatch caused by the false light cue. Alternatively, nocturnal begging may serve a social function by signalling individual needs, mediating sibling competition, and influencing parental allocation, with its expression shaped by brood size, nestling sex, and nutritional needs (Leonard, 2000; Price et al., 1996; Romano et al., 2012). Nestlings exposed to ALAN may remain more active at night instead of resting, leading to sleep disruptions. Compared to DARK, nestlings in ALAN exhibited more restlessness at night, moving around the nest cup and wiggling (Fig. S3). It is yet to be determined if the night-time begging results from increased hunger via higher ghrelin secretion, the appetite-regulating hormone, due to a shift in the circadian clock. Alternatively, it may simply contribute to increased energy expenditure, potentially leading to poorer body condition and impaired survival. Reduced melatonin and sleep disruption may also drive this increased nocturnal activity. However, in nestlings, the restlessness manifested as increased begging (Fig. 2) makes it difficult to separate true energetic need from mere wakefulness. Indeed, excessive begging brings about various costs, for example, impaired growth, body mass, immunity and reproductive success (Fresneau & Müller, 2019; Kilner, 2001; Redondo et al., 2016). Although it is possible to compensate for daytime mass loss at night by losing less mass (Redondo et al., 2016), ALAN-induced sleep disruption may undermine this recovery, by amplifying the physiological costs of begging. Importantly, our study also suggests such costs, as restless, begging nestlings exposed to ALAN delayed fledging and showed reduced fledging success compared to controls (Fig. 5c,d). Contrary to our expectations, ALAN nestlings did not decrease the frequency or duration of daytime begging (Fig. 2a,b), which may reflect stimulation by the presence of parents, consistent with the positive phenotypic covariation between begging and parental provisioning (Fresneau & Müller, 2019).

Parents extended their activity period, by stopping their visits later in the evening and starting earlier in the morning (Fig. 3, Table 2), in accordance with previous studies investigating activity onset and offset in the wild (Beaugeard et al., 2024; Stracey et al., 2014; Wang et al., 2021). Although the flycatchers were free-ranging, nestboxes were placed deep in wooded areas, and light pollution in rural Gotland was low. This minimized adult exposure to ALAN outside the nestboxes. Thus, the extension of activity likely resulted directly from in-box illumination, particularly for brooding females, and indirectly from socially mediated cascades via parent-offspring (Fresneau & Müller, 2019) and male-female communication (Baldan & van Loon, 2022; van Rooij & Griffith, 2013). Such disruption of natural daily rhythms through behavioural compatibility may elevate the energetic costs of parental care (de Jong et al., 2016; Spoelstra et al., 2018). On the other hand, from the nestling’s perspective, ALAN appeared to enhance the feeding opportunity window by improving visibility inside the nestbox after sunset. Nevertheless, this extension of daily parental care was traded off with the reduced hourly feeding rate by both parents (Table 3, Fig. 4a,b), consistently with previous evidence (Injaian et al., 2021; Titulaer et al., 2012; Welbers et al., 2017). While daytime begging frequency did not differ across treatments (Fig. 1a), females were generally more responsive to begging (Table 3), with those in ALAN nests showing stronger responses than females in DARK nests. However, this pattern should be interpreted cautiously, as begging ranges did not fully overlap between treatments (Fig. 4c). The clearer feeding-begging relationship under ALAN likely reflects more uniform begging levels, while the broader range in DARK may have diluted the response. Male feeding appeared less sensitive to begging cues, increasing instead with brood size (Fig. 4d) and showing a tendency to rise with the proportion of male offspring (Table 3). As such, fathers’ investment was skewed towards high-quality and male-biased broods. These findings collectively indicate higher parental investment by females, who sustained feeding effort despite a prolonged active period under ALAN (Fig. S6a). This aligns with prior work in great tits (Titulaer et al., 2012), which also reported increased female care under ALAN, possibly due to greater exposure while in the nest. Females may remain longer to perform nest sanitation, increasing their exposure to both ALAN and begging stimuli (Lucass et al., 2016). On day 8, some females brooded overnight, with more doing so in DARK nests (38%) than in ALAN nests (22%). Although this difference was not significant (*Χ^2^* = 1.12, df = 1, p = 0.290), it suggests a potential female behavioural adjustment to light.

Contrary to our hypothesis, but in accordance with some prior evidence (Ziegler et al., 2021), we did not observe an unequivocal effect of ALAN on nestling growth parameters (Table S6). We, however, noted a tendency for enhanced structural wing growth under ALAN (Table S6a), explainable by prolonged feeding window by both parents and enhanced response to begging by females. However, the lack of differences in body mass could stem from the reduced feeding rate by parents and increased net energy expenditure of ALAN nestlings that begged at night (Raap, Casasole, et al., 2016) (Fig. 1a,b). Additionally, there was no difference in the body condition of nestlings at day 12 and the number of fledged offspring (Table 4a,b). Similar findings have been reported in previous studies, where ALAN either had no effect (Grunst et al., 2020) or increased fledging success (Wang et al., 2021). In our study, body condition and fledging number have only been positively influenced by feeding rates, independently of ALAN (Fig. 5a,b). Despite no overall difference in fledging number, ALAN significantly reduced the fledging rate from day 8 onward, indicating that light exposure affected survival during the late nestling phase (Table 4c). Early mortality likely reflected intrinsic factors such as initial viability or early parental investment (Martin, 1995), whereas later mortality possibly relied more on physiological sensitivity to light. Although circadian melatonin biosynthesis begins in birds during embryonic development, it stabilises later on, when nestlings become increasingly responsive to light-dark cues (Zeman & Herichová, 2011). This marks a critical window in which ALAN may disrupt hormonal regulation, behaviour, and parent-offspring interactions (Grunst & Grunst, 2023). Our findings indicate that nests with higher feeding rates under ALAN showed improved fledging success (Table 4c, Fig. 5c), highlighting that compensatory parental investment may mitigate ALAN’s physiological costs during this vulnerable period. Most remarkably, we show that ALAN delayed fledging (Table 4d, Fig. 5d). While from the perspective of nestlings it is potentially beneficial to develop more before fledging, extended nesting times are also associated with increased mortality due to predation (Martin, 1995) or, particularly in the case of time-pressured migrating species, with delays in preparations for migration (Bani Assadi & Fraser, 2021). For parents, the combination of longer in-nest feeding period and extended daily activity imposes additional reproductive costs, potentially reducing future breeding opportunities and overall fitness (Gustafsson & Sutherland, 1988). Such increased demands have been shown to limit the energy available for moulting, resulting in lower-quality feathers that compromised flight performance and raised thermoregulatory costs (Dawson et al., 2000; Nilsson & Svensson, 1996). For long-distance migratory birds like the collared flycatcher, the timing of moulting and the quality of their plumage are critical for successful migration (Barta et al., 2008).

In conclusion, while ALAN directly impacted primarily nestlings and, to a lesser extent, brooding females, its behavioural effects extended beyond these individuals. Social interactions and intra-family communication appeared to mediate broader consequences, influencing the circadian behaviour of entire bird families, leading to extended nestling periods and reduced fledging success. This suggests that ALAN may disrupt avian family units through direct exposure and indirect, socially transmitted effects, demonstrating the need to consider group-level dynamics in assessing ecological responses to anthropogenic light pollution.

## Ethics approval

All methods and sampling detailed in this manuscript were performed according to Swedish regulations concerning work with wild populations and under the approval and permission of Jordbruksverket no. 9164-2021.

## Acknowledgements

We thank members of the Gotland fieldwork team and numerous field assistants, particularly Wioleta Oleś, Martyna Cendrowska, Junchen Deng and Ninon Gorget, for their contribution to data collection. We also thank Marek Sudyka for preparing light sets and Arduino programming, Wioleta Oleś for performing molecular sexing of the nestlings and Paweł Cembrzyński for helping with the graphics.

## Funding

The study was funded by the NCN grant, SONATA no. 2019/35/D/NZ8/00889 awarded to JS. We additionally thank the NCN grant no. 2020/39/B/NZ8/01157 for supporting the fieldwork of MC and general bird monitoring.

## Author contributions

JC and JS conceived and designed the study. JC, SDP, IDL, MC and JS performed the fieldwork. JC analysed the videos. JC and JS performed statistical analyses. LG provided logistical support in the field. JC and JS wrote the manuscript with input from all authors. All authors approved the final version of the manuscript.

## Competing interests

The authors declare no competing interests.

## Data and materials availability

The R code used in this study and raw data can be found at https://doi.org/10.5281/zenodo.15608337 (Champenois et al., 2025).

## Notes

https://zenodo.org/records/15608337

### Competing Interest Statement

The authors have declared no competing interest.

