## Supplementary Material for "Awake and hungry: artificial light at night disrupts behaviour and reproductive ecology in a wild migratory bird"

**Affiliations**:

### Light sets

Each light set consisted of a plate with Arduino and a clock with a battery plus resistor that generated a power load every 10 seconds to keep the power bank active, a cable which was connected to a power bank and a LED light bulb on a wire (Fig. S1). The sets were programmed to switch the light on at 19h00 and switch it off at 06h00. The LED white light diode was placed inside the nestbox, under the lid using a bendable metallic plate. We adjusted the diode illuminance to produce 1.5 lx at the distance of 17 cm from the light source, that is 10 cm above the nestbox bottom (Fig. 1b) to standardize the light intensity perceived at the level of developing nestlings. With the advancement of the season the nest cup is being compressed, and the average nest height in the studied population decreases from 9.0 cm (SD ± 2.58) to 5.2 cm (SD ± 2.49), measured from the nestbox bottom in 18 nests at day 2 of nestling's life and 98 nests at day 8 of nestling's life, respectively. The Arduino boards and batteries were stored in a sealed plastic box outside the nestbox, along with a power bank for the camera during recording time. For control sets, there was only a dummy LED diode and a plastic box for the camera's power banks to equalise the conditions between experimental groups. During each visit, the power banks were checked and replaced if needed.

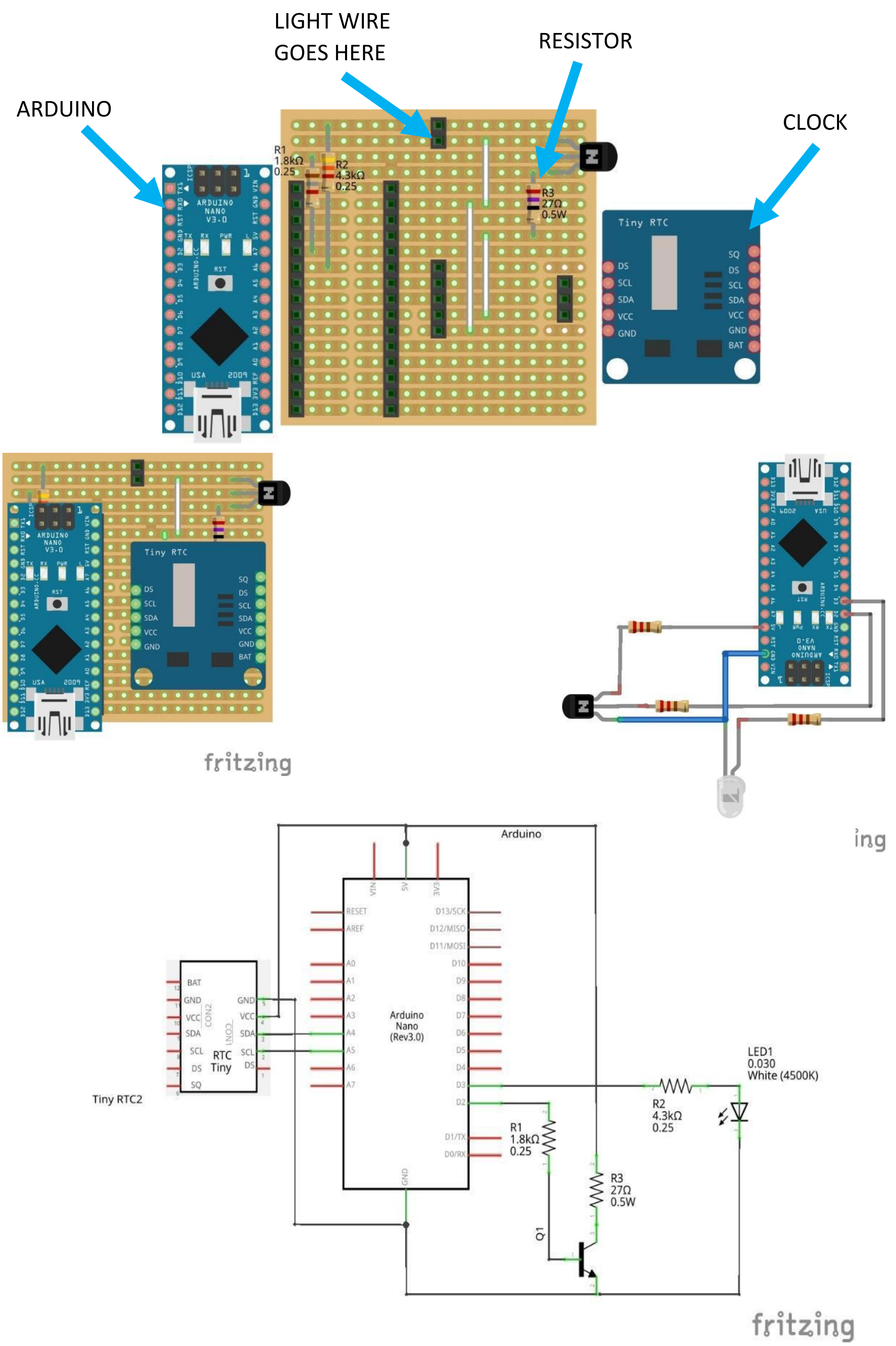

*Figure S1 - Light set scheme*

### Cameras

To record the behaviour inside the nest, small spy cameras (8 model KA9 by kamREC in 2022 and 8 additional model A10 by NOBITECH SC in 2023) (Fig. S2) were magnetically fixed to a metallic plate next to the diode and secured with tape. The cameras had a diameter of 4.5 cm and a height of 2.5 cm. They could record in dark conditions using the infrared spectrum (940nm). The recordings were saved in MP4 format on micro-SD cards.

In the field, cameras were connected through WIFI to a smartphone using the "HIDVCAM" application to adjust the settings in order to ensure that the whole nest was visible on the screen (Fig. S2). The date and time from the camera were automatically synchronised with the smartphone's clock and displayed on the recordings. The camera recorded on motion detection to save battery but with the sensitivity set to "high" to record all activity within the nest. The night vision was enabled and the light indicators were disabled.

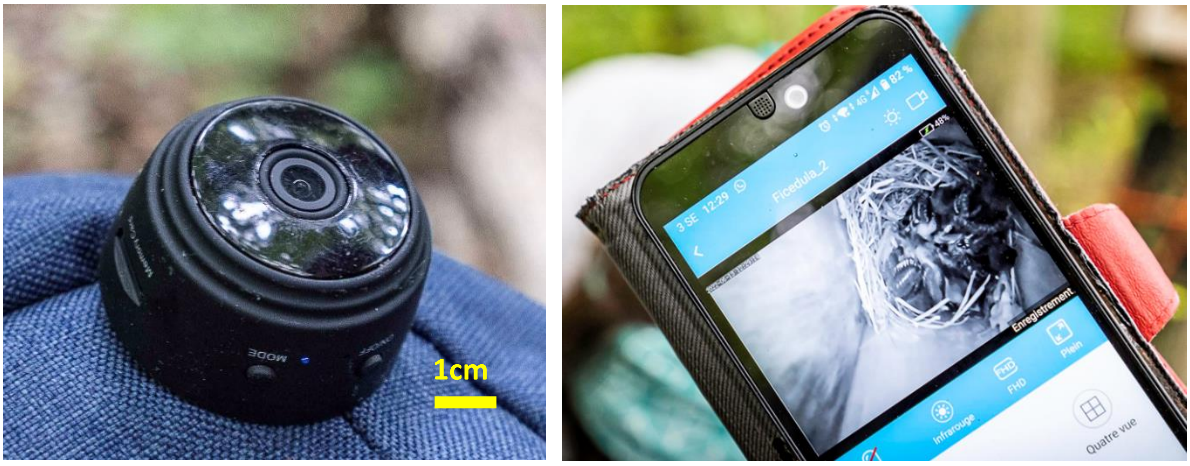

***Figure S2 - Type of camera used for recordings, live feed of camera connected to a smartphone***

### Fieldwork on Gotland

Nestboxes were inspected every four days from the end of April to record all breeding attempts. Aside from collared flycatchers, they can be primarily occupied by blue tits (*Cyanistes cearuleus*) or great tits (*Parus major*), which begin breeding circa 2 weeks earlier than the collared flycatchers and can compete for the breeding space. After identifying an active flycatcher nest from the appearance of nesting material, we started individual nest monitoring. We recorded lay date (LD), clutch size (CS) and incubation start (IS). The incubation was determined via two ways: first, by checking if the eggs were warm, then egg candling was used to more accurately determine the start of embryo development (Ojanen & Orell, 1978), and minimize the visits to the nests for hatching checks. Two nests with the same hatching date (HD ± 1 day) and the same number of hatchlings (brood size ±1) were paired, and randomly assigned to either experimental (ALAN) or control (DARK) group.

Two days after hatching, we installed the light sets and began the experimentation. The experimental timeline is shown in detail in Fig. 1c. Cameras were installed on average at 14:00 at HD + 8 (that occasion was also used to uniquely ring the nestlings) and removed on average at 15:00 the next day. During each visit, additional biometric data were collected (Table S1).

***Table S1 - Description of the measured biometric parameters***

| **Parameter** | **Day of measurement** | **Description** |
|---|---|---|
| Body mass | Day 2, 8, 9 and 12 | Body mass (grams) of the nestling |
| Wing chord length | Day 8, 9 and 12 | Distance of the closed wing from the foremost extremity of the carpus to the tip of the longest primary feather (mm) |
| Tarsus | Day 12 | Length of the tarsometatarsal bone (mm) |
| Ectoparasites | Day 12 | Presence or absence of ectoparasites on each nestling, i.e., ticks, fleas, lice, blowfly larvae, *Hippoboscidae* |

Starting from HD + 15, the nestboxes were checked for fledging every two days. To minimise disturbance and not to induce fledging, only if the parents could not be seen going to the nest when observing for 10 minutes from a distance, or the nestlings could not be heard, was the box opened. We determined the exact fledging day and hour (data not shown) by analysing the data from temperature loggers placed inside the nestbox during day 12 measurements (Fig. 1c) and set to hourly recordings. When fledging was confirmed, the light sets were removed and dead nestlings (if any) were identified by their ring number to determine fledging success. A total of 45 nests were recorded and analysed in 2022 and 2023. 21 nests in the 2024 season were added for the analyses of first and last parental visit (see Table 3).

***Table S2 - Sample sizes showing the number of nests per year and per experimental group included in the study***

| **YEAR** | **DARK** | **ALAN** | **TOTAL** |
|---|---|---|---|
| 2022 | 8 | 6 | 14 |
| 2023 | 16 | 15 | 31 |
| 2024* | 13 | 8 | 21 |
| TOTAL | 24/37* | 21/29* | 45/66* |

*2024 included only for the analyses of last and first parental visit

### Sexing of the nestlings

DNA for sexing was extracted from blood samples collected in 98% ethanol immediately after sampling via brachial vein puncture at day 12 after hatching (ca 30 ul per nestling). For some samples in 2023 and all samples in 2022 the blood was collected in RNAlater. The extraction was performed with Chelex (Walsh et al., 1991) and PCRs were performed with standard primers P2 and P8 (Griffiths et al., 1998) with the following specific conditions:

PCR conditions (10 ul)

| | | ul/sample |
|---|---|---|
| Buffer (green) | 10x | 1 |
| MgCl2 | 25 mM | 0.6 |
| P2 | 10 uM | 0.5 |
| P8 | 10 uM | 0.5 |
| Dream Taq | 5U/u | 0.1 |
| dNTP mix | 2 mM | 0.1 |
| H2O | | 6.2 |
| DNA | | 1 |

Program (35 cycles)

| | |
|---|---|
| 94°C | 2 min |
| 94°C | 30 s |
| 50°C | 45 s |
| 72°C | 1 min |
| 72°C | 10 min |

The PCR product was then assessed for either two bands (female ZW chromosomes) or one band (ZZ) by electrophoresis on agarose gel (1.5-2%) at 180 V for 120 min. Information on individual sex was used to calculate the proportion of male offspring, which was the number of males over the number of nestlings sexed within the brood. It was included in statistical modelling to account for sex-specific nutritional requirements of nestlings, which can influence parental provisioning and fitness outcomes (Sheldon et al., 1998). We only included broods in which we successfully sexed at least 50% of the nestlings alive at day 8; this led to the exclusion of two DARK nests in all analyses using this factor as a predictor.

### Video analysis

After retrieving SD cards from the field, recordings were transferred to a computer and backed up with two separate copies on hard drives. Due to the motion detection settings, many short videos were recorded as the camera would stop in the absence of movement. Hence, the videos were merged using Shotcut, a free video editing software to create a collated video of all the activity recorded within the hour of day or night.

An ethogram (Table S3) was established by precisely watching one hour of night and one hour of day activity in a random pair of nests (one in DARK and one in ALAN). While we recorded ca. 24h of activity per nest, we decided to reduce the detailed analysis to just two hours because of the number and duration of the recordings. To test whether one hour is representative for overall nightly activity within the nest, we initially analysed the activity for two hours per night, from 23:00 to 0:00 and from 01:00 to 02:00 in 12 nests (five experimental and seven control nests recorded in 2022). Since the begging behaviour we focused on was highly correlated between the two hours ($r = 0.97$, $N = 12$, $p < 0.0001$), only the hour between 01:00 and 02:00 was analysed in the rest of the nests and used for the analyses.

The daily behaviour of the nestlings and parents was analysed between 09:00 and 10:00 invariably in all nests. To choose the time most representative for average daily activity behaviour, we examined the length of recordings per each hour between 08:00 and 12:00 of 6 random nests (three experimental and three control nests). Since the cameras were operating in motion detection mode, the recording length was a good approximation of general hourly activity. Activity between 9:00 and 10:00 was moderate, relative to peaks and lows at other times.

The behavioural data was obtained from the videos using BORIS (Behavioural Observation Research Interactive Software) (Friard & Gamba, 2016). During the screening, the only information available was: the number of nestlings, the ID of the nest, the day and time, so the screening was performed blindly regarding the treatment to reduce the experimenter bias. Night videos were screened first since our main focus is the begging behaviour of nestlings at night. Screening day videos took longer due the overlapping presence of different focal subjects (the nestlings, the female and the male). The nests were analysed in a random order, and it was impossible to determine if the nest belonged to the control (DARK) or the experimental (ALAN) group.

***Focal subjects***

During the analysis, each behaviour was attributed to one subject: the male, the female or the nestlings. The breeding plumage of adult male flycatchers is black on the head, back, wings and tail, with a white collar, large white patches on the forehead and wings, and a white belly. They are easily distinguishable from females, who are grey brown above and white below. In the rare occurrences the adult could not be identified with certainty (i.e., fed the nestlings from the nest opening without entering the frame long enough to capture a clear image) or in the unique event of two male occurring in the frame at the same time, they were just referred to as a "parent" and were removed from the analysis. Adults' behaviours encompassed: feeding, entering the nest and cleaning (Table S3). Since the nestlings were not marked, it was not possible to record individual behaviours. Thus, nestlings' behaviour was recorded if at least one nestling displayed a behaviour. Nestling behaviours encompassed begging, moving, scratching, and wiggling (Table S3). In the seven cases where the female stayed in the nest at night, it was difficult to determine if the movements observed originated from the female or the nestling under her. Since the female was the subject visible on the screen, the behaviours were recorded with her as a subject hence, and not included in the night analyses of nestlings' behaviour.

***Behaviour types***

BORIS allows to record two types of behaviour, a point event and a state event. The main difference is that the former is a single occurrence and the latter is a continuous event with the duration expressed in seconds. The behaviours, their types, associated subject and description are presented in Table S3. Across all nests we recorded 32,100 behaviours in nestlings (17,457 at night and 14,643 during the day) and 3,709 in parents (401 at night and 3,308 during the day). 17 nestling behaviours were assigned as other (mostly wing flapping) and were removed from the subsequent analysis (Fig. S3). In 28 cases, the sex of the parent could not be assigned and these were removed. For the comparative analysis of the full set of circadian parental behaviours according to manipulation (Fig. S4), "enter nest" was removed as this state event informed when the parent appeared on the recording and encompassed all other types of behaviour. Night behaviours were only recorded in females that spent the night in the nests: wiggling (156), sleeping (128) in seven nests, scratching (62), moving around (16), other (e.g., pecking and rearranging the nest, 38).

***Table S3 - Description of recorded behaviours***

| Behaviour | Subject | Event type | Description |
|---|---|---|---|
| Beg | Nestlings | State | Start: At least one nestling raises its head, opening its beak. Stop: all nestlings have closed their beaks |
| Move around | Nestlings, Female, Male | Point | The subject is shifting position within the nest cup |
| Wiggle | Nestlings, Female, Male | Point | The subject moves, but its position within the nest cup does not change |
| Scratch | Nestlings, Female | Point | The individual rubs its body with its beak or leg |
| Feed | Female, Male | Point | The subject puts an item in the beak of one of the nestlings |
| Clean | Female, Male | Point | The subject picks up faeces from the nest |
| Enter nest | Female, Male | State | Start: the focal subject appears in the frame. Stop: the subject leaves the frame |
| Sleep | Female | State | Female sleeps in 7 nests, assuming the sleeping position (head under the wing) (Raap et al., 2016) |
| Sit | Female | State | Female sits without moving |
| Allofeeding | Male | Point | Male feeds the female on the nest |

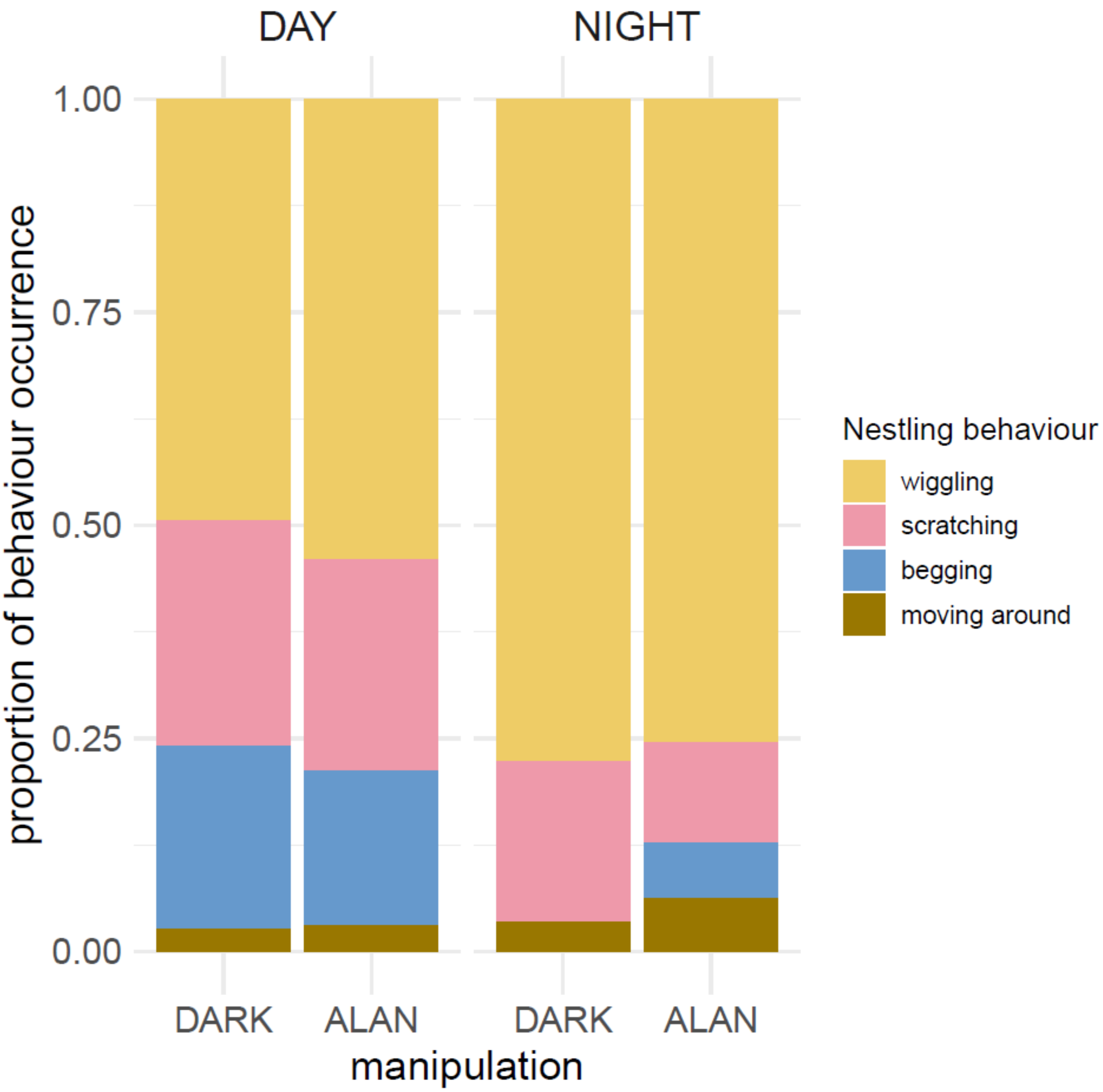

***Figure S3 - Proportion of the full set of circadian nestling behaviours according to manipulation*** for day and nighttime. 32,083 (14,626 day and 17,457 night) behaviours, differed between DARK and ALAN during the day ($X^2 = 41.32$, df = 3, $p < 0.0001$) and at night ($X^2 = 717.76$, df = 3, $p < 0.0001$). In terms of counts of behaviours other than begging, compared to DARK, nestlings in ALAN were less restless during the day: less scratching ($X^2 = 79.29$, df = 1, $p < 0.0001$) and less wiggling ($X^2 = 34.65$, df = 1, $p < 0.0001$) with no difference in moving around ($X^2 = 0.63$, df = 1, $p = 0.426$) and more restless during the night: more moving around ($X^2 = 116.30$, df = 1, $p < 0.0001$) and more wiggling ($X^2 = 34.65$, df = 1, $p < 0.0001$), however they were scratching less ($X^2 = 51.66$, df = 1, $p < 0.0001$).

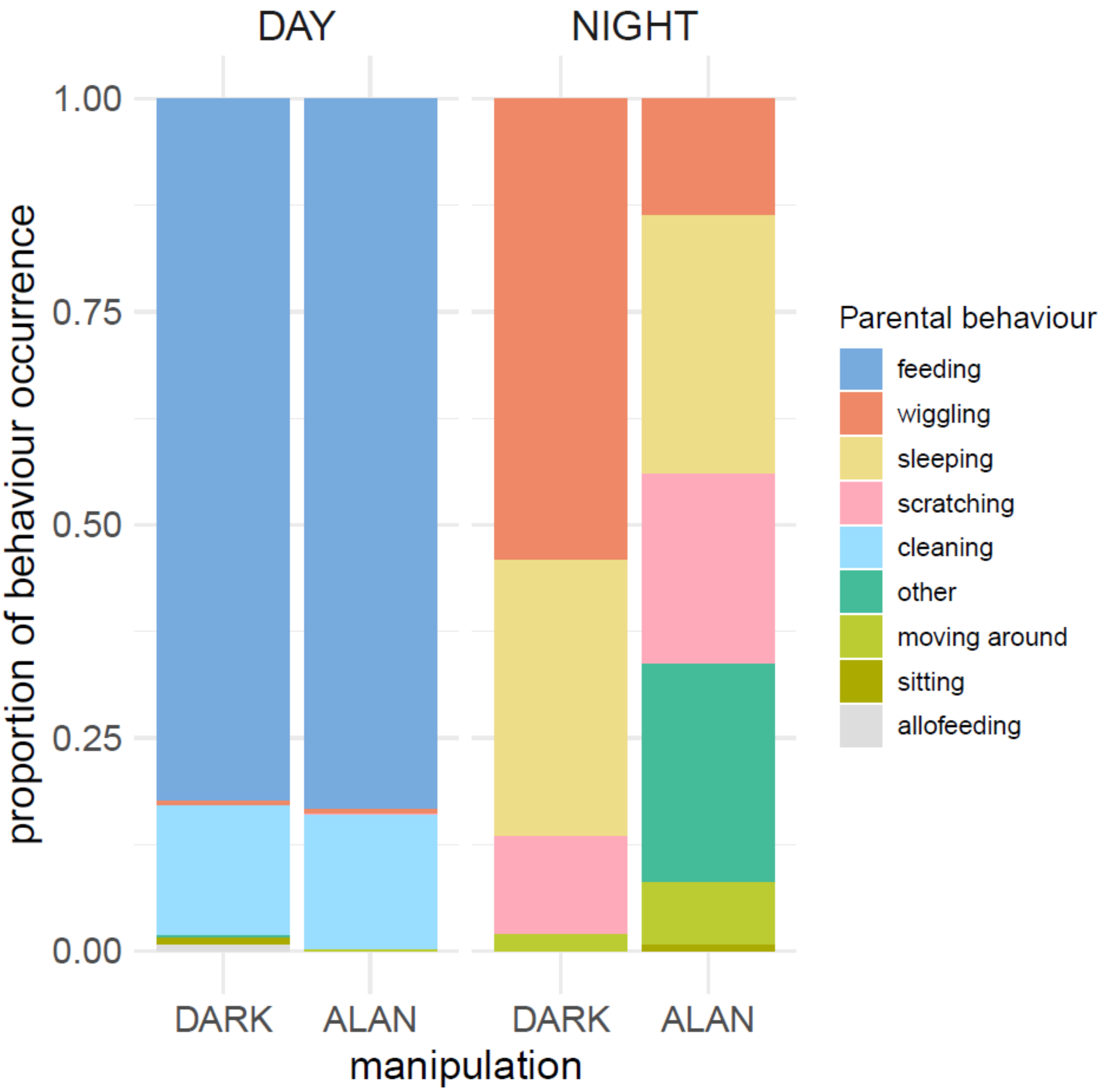

***Figure S4 – Proportion of the full set of circadian parental behaviours according to manipulation*** for day and nighttime. 2,182 (1,782 day and 400 night) behaviours, differed between DARK and ALAN during the day ($X^2$ = 14.90, df = 7, p = 0.037) and at night $X^2$ = 119.59, df = 5, p < 0.0001).

We then focused on parameters related to begging and feeding behaviours. In nestlings, we quantified per nest: Total Begging Count (number of begging events summarised for night and day), Begging Day (hourly number of begging events during the day), Begging Night (hourly number of begging events during the night), Begging Duration (average duration of a begging event). In parents we quantified per nest: Feeding Count (total hourly number of feeding events), Male Feeding Count (hourly number of feeding events by male), Female Feeding Count (hourly number of feeding events by female), Parent Visit Duration (average duration of parental visit during the day), Entering Nest Count (hourly number of parental visit events during the day). We observed strong correlations among some of the parameters (Fig. S5). For example, Entering Nest Count and Feeding Count correlation: $r = 0.873$, $t = 11.63$, $df = 42$, $p < 0.0001$.

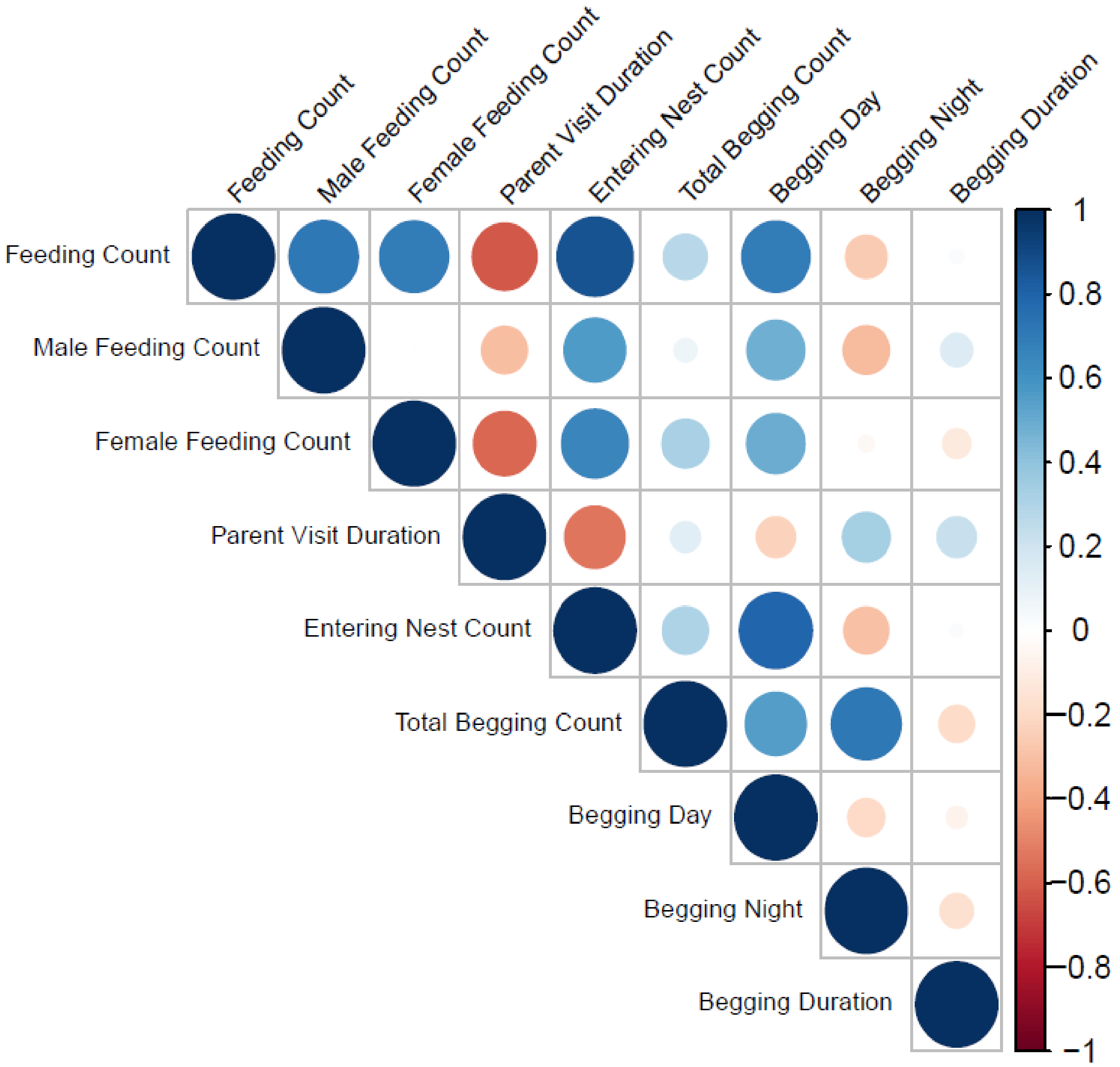

***Figure S5 - Correlation heatmap of the behavioural parameters related to begging and feeding*** video-recorded per nest. Blue denotes positive correlations, and red negative. The dot size and colour intensity denote correlation strength, with darker colours and larger dots indicating stronger correlations.

***Last and first parental visit***

The parental visit times were established based on specific criteria. The last visit time was pinpointed as the time the parent departed from the nestbox before the evening's decline in activity on the installation day (HD + 8). This was assessed separately for males and females. For the seven instances where the female remained in the nest, it was identified as the moment prior to her settling in for the night. The time of last visit was converted to minutes after sunset for the subsequent analysis.

Conversely, the first visit time was determined by the initial appearance of the parent in the frame, the morning after the camera was set up (HD + 9). Again, this was analysed separately for males and females, with the exception of the seven cases where the female remained in the nest; in those situations, it was recognized as the first reappearance after her initial departure. The time of first visit was converted to minutes after sunrise for the subsequent analysis.

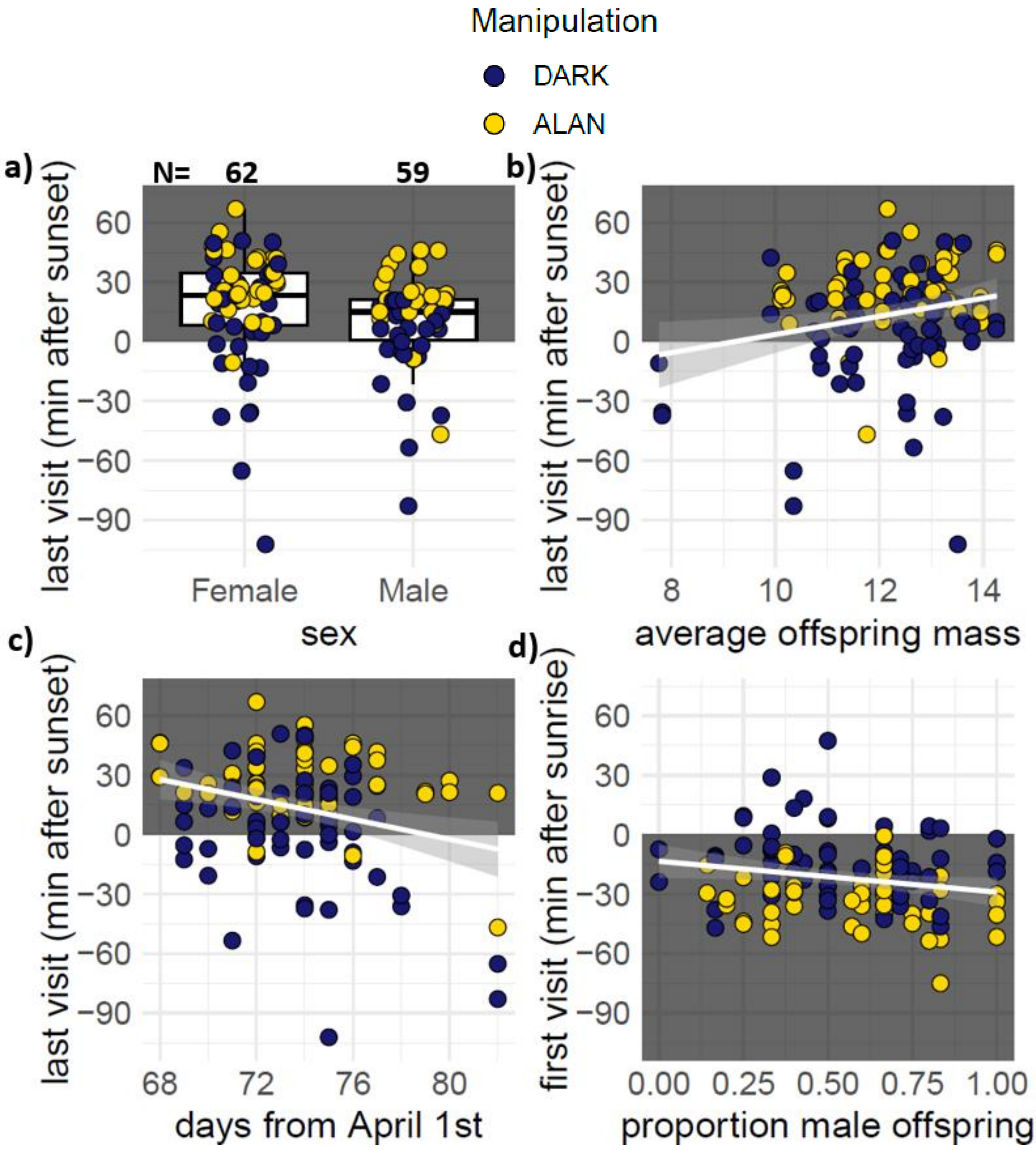

***Figure S6 – Timing of last parental visit to the nest*** according to a) parental sex, b) average offspring mass on day 8th, c) timing in the reproductive season and d) the timing of first parental visit to the nest according to a proportion of male offspring in the brood (increasing values denote more male offspring) under ALAN (yellow) and DARK (dark blue) conditions. A white background indicates daytime (before sunset and after sunrise), while nighttime (after sunset and before sunrise) is indicated by a grey background. In a), the boxes represent the median with IQR, and the whiskers 1.5 IQR; in b), c) and d), the white lines show the fitted linear relationships with shaded standard error bands. Plotted on raw values.

***Table S4a - Pairwise comparisons of Estimated Marginal Means for manipulation and time interaction for Model 1a on circadian begging count*** from Table 1a. Significant differences (P < 0.05) are indicated in bold

| **Contrast** | **Estimate ± SE** | **z.ratio** | **p.value** |
|---|---|---|---|
| DARK Day - ALAN Day | 0.23 ± 0.10 | 2.39 | 0.078 |
| DARK Day - DARK Night | 4.64 ± 0.46 | 10.06 | **< 0.0001** |
| DARK Day - ALAN Night | 0.96 ± 0.12 | 7.86 | **< 0.0001** |
| ALAN Day - DARK Night | 4.41 ± 0.46 | 9.52 | **< 0.0001** |
| ALAN Day - ALAN Night | 0.72 ± 0.13 | 5.63 | **< 0.0001** |
| DARK Night - ALAN Night | -3.68 ± 0.47 | -7.87 | **< 0.0001** |

***Table S4b - Pairwise comparisons of Estimated Marginal Means for manipulation and time interaction for Model 1b on circadian begging duration*** from Table 1b. Significant differences (P < 0.05) are indicated in bold

| **Contrast** | **Estimate ± SE** | **z.ratio** | **p.value** |
|---|---|---|---|
| DARK Day - ALAN Day | 0.01 ± 0.04 | 0.30 | 0.990 |
| DARK Day - DARK Night | 1.42 ± 0.28 | 4.99 | **< 0.0001** |
| DARK Day - ALAN Night | 0.56 ± 0.05 | 11.85 | **< 0.0001** |
| ALAN Day - DARK Night | 1.40 ± 0.28 | 4.92 | **< 0.0001** |
| ALAN Day - ALAN Night | 0.54 ± 0.04 | 13.51 | **< 0.0001** |
| DARK Night - ALAN Night | -0.85 ± 0.29 | -2.99 | **0.015** |

***Table S5 - Model examining variation in begging duration (natural log-transformed for normality of the model residual distribution) in response to the average duration of parental visit*** during the day. Predictor variables were selected based on model averaging across all models with ΔAICc <2, where Manipulation:ParentVisitDuration was always retained as a *fixed* factor of interest. Significant differences ($P < 0.05$) are indicated in bold. Marginal ($R^2$m) and conditional ($R^2$c) R-squared are shown. Since the focal interaction Manipulation:ParentVisitDuration was non-significant (0.011 ± 0.007, t = 1.34, p = 0.188), we removed it for the correctness of the main factor interpretation.

| **Model 2: begging duration day,** LMM gaussian | | | | |
|---|---|---|---|---|
| **N = 42 nests** | | | | |
| | **Estimate** | **SE** | **t** | **p** |
| **(Intercept)** | 1.34 | 0.18 | 7.97 | **<0.0001** |
| **Manipulation (ALAN)** | -0.02 | 0.05 | -0.29 | 0.771 |
| **ParentVisitDuration** | 0.007 | 0.004 | 1.84 | 0.074 |
| **BroodSizeD8** | 0.09 | 0.02 | 3.90 | **0.0004** |
| ***Random effects*** | ***Variance*** | | | |
| ***Pair*** | *0.001* | | | |
| ***Site*** | *0 (removed)* | | | |
| **$R^2$m / $R^2$c** | 0.280 / 0.295 | | | |

***Table S6 - Models examining variation in a) average structural size (wing) gain and b) body mass gain in response to parental hourly feeding count*** between days 8 and 9 of age when the video recordings took place. Predictor variables were selected based on model averaging across all models with ΔAICc <2, where Manipulation:FeedingCount was always retained as a *fixed* factor of interest. The best model is presented for mass gain (since just one model's ΔAICc <2). Significant differences ($P < 0.05$) are indicated in bold. Marginal ($R^2$m) and conditional ($R^2$c) R-squared are shown. Since the focal interaction Manipulation:FeedingCount in the wing gain model (0.006 ± 0.020, t = 0.30 p = 0.771) and mass gain model (0.02 ± 0.01, t = 1.74 p = 0.095) were non-significant, we removed those for the correctness of main factor interpretation in the models.

| **a. Model 5: wing size gain, LMM gaussian** | | | | |
|---|---|---|---|---|
| **N = 42 nests** | | | | |
| | **Estimate** | **SE** | **t** | **p** |
| **(Intercept)** | 8.74 | 1.10 | 7.92 | **<0.0001** |
| **Manipulation (ALAN)** | 0.43 | 0.24 | 1.80 | 0.094 |
| **FeedingCount** | 0.04 | 0.02 | 2.63 | **0.016** |
| **BroodSizeD8** | -0.89 | 0.16 | -5.69 | **<0.0001** |
| **Year (2023)** | -1.31 | 0.85 | -1.55 | 0.155 |
| ***Random effects*** | ***Variance*** | | | |
| ***Pair*** | *1.543* | | | |
| ***Site*** | *0.063* | | | |
| ***HoursBetweenMeasurements*** | *0.068* | | | |
| ***ObserverD8*** | *0 (removed)* | | | |
| ***ObserverD9*** | *0.418* | | | |
| **$R^2$m / $R^2$c** | 0.404/ 0.945 | | | |

| **b. Model 6: body mass gain, LMM gaussian** | | | | |
|---|---|---|---|---|
| **N = 42 nests** | | | | |
| | **Estimate** | **SE** | **t** | **p** |
| **(Intercept)** | 1.04 | 0.33 | 3.15 | **0.003** |
| **Manipulation (ALAN)** | 0.09 | 0.16 | 0.54 | 0.593 |
| **FeedingCount** | -0.001 | 0.00[illegible] | -0.12 | 0.902 |
| ***Random effects*** | ***Variance*** | | | |
| ***Pair*** | *0.143* | | | |
| ***Site*** | *0 (removed)* | | | |
| ***HoursBetweenMeasurements*** | *0 (removed)* | | | |
| **$R^2$m / $R^2$c** | 0.007/ 0.412 | | | |

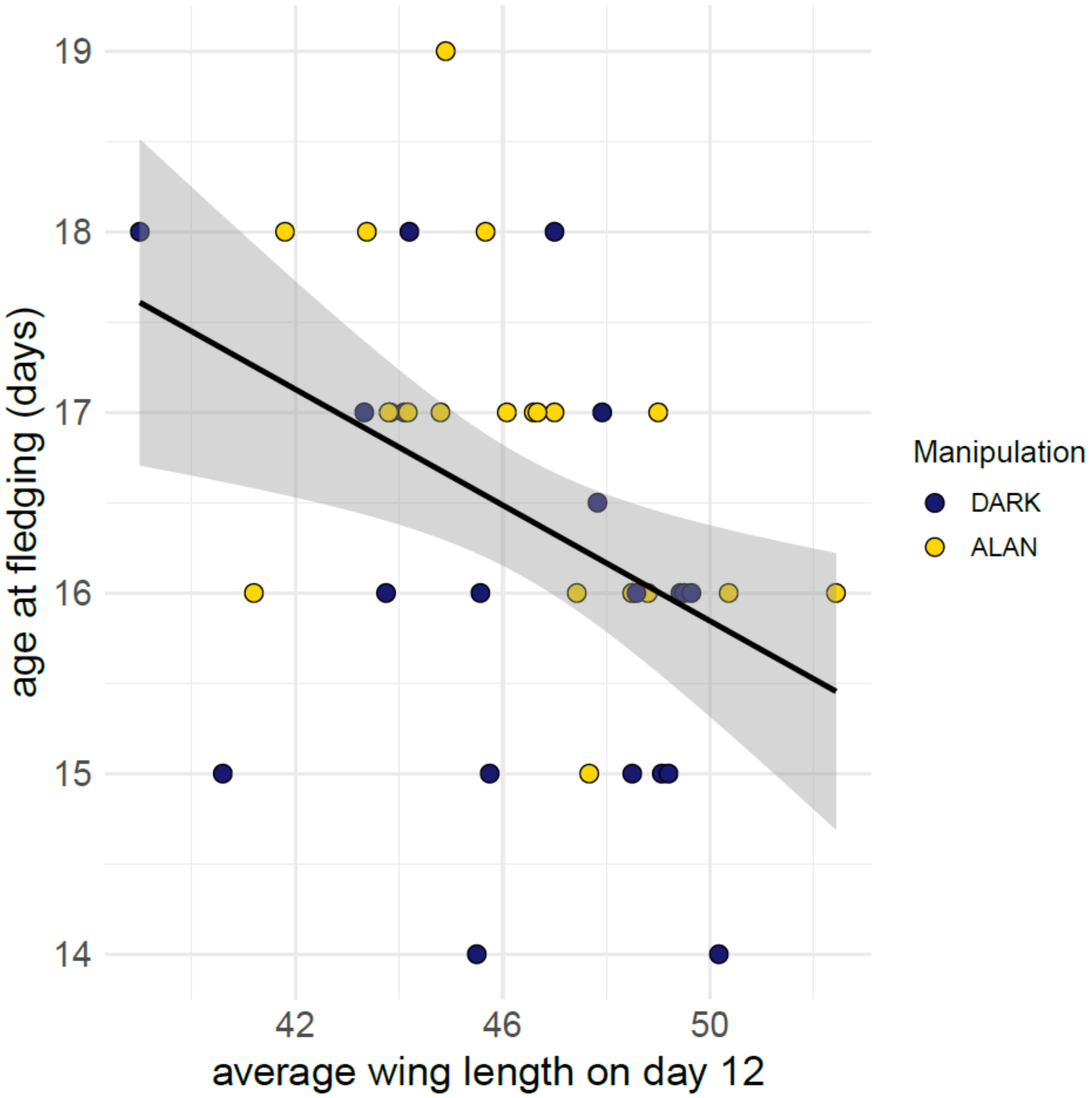

***Figure S7 – Age at fledging in relation to average wing length on day 12*** under ALAN (yellow) and DARK (dark blue) conditions. The line shows the fitted linear relationship with shaded standard error bands. Plotted on raw values.
